# The Vacc-SeqQC Project: Benchmarking RNA-Seq for Clinical Vaccine Studies

**DOI:** 10.1101/2022.06.23.497396

**Authors:** Johannes B. Goll, Steven E. Bosinger, Travis L. Jensen, Hasse Walum, Tyler Grimes, Gregory D. Tharp, Muktha S Natrajan, Azra Blazevic, Rich Head, Casey E. Gelber, Nirav B. Patel, Patrick Sanz, Nadine G. Rouphael, Evan J. Anderson, Mark J. Mulligan, Daniel F Hoft

**Author notes:** To whom correspondence should be addressed: Daniel F Hoft, Daniel.Ho, Mark J Mulligan. Contributed equally (Johannes B. Goll, Steven E. Bosinger, Mark J. Mulligan, Daniel F Hoft).

## Abstract

Over the last decade, the field of systems vaccinology has emerged, in which high throughput transcriptomics and other omics assays are used to probe changes of the innate and adaptive immune system in response to vaccination. RNA-Seq technology has matured in recent years and is now widely deployed for transcriptional analysis of clinical specimens. The goal of this study was to benchmark technical parameters of RNA-Seq in the context of a multi-site, double-blind randomized clinical trial using primary patient samples. We collected longitudinal peripheral blood mononuclear cell (PBMC) samples from 10 subjects after vaccination with a live attenuated *Francisella tularensis* vaccine and performed RNA-Seq at two different sites using aliquots from the same sample to generate two large-scale replicate datasets. We evaluated the impact of: (i) filtering lowly-expressed genes, (ii) using external RNA controls, (iii) fold change and false discovery rate (FDR) filtering, (iv) read length, and (v) sequencing depth on differential expressed genes (DEGs) concordance between replicate datasets. Using synthetic mRNA spike-ins, we developed a method for empirically establishing minimal read-count thresholds for maintaining fold change accuracy on a per-experiment basis. We defined a reference PBMC transcriptome by pooling sequence data, and established the impact of read depth and gene filtering on transcript representation. Lastly, we modeled statistical power to detect DE genes for a range of sample sizes, effect sizes, and coverage depths. The results from this study provide RNA sequencing benchmarks and guidelines for planning future similar vaccine studies.

## Introduction

Since 2008, high-throughput technologies, primarily transcriptomics, have been used to characterize the *in vivo* response to clinical vaccination (1, 2). This approach, referred to as “Systems Vaccinology” has been employed to investigate the molecular mechanisms regulating vaccine activity (3), to identify correlates that predict antibody titer, breadth, or persistence (4-7), and to predict responses to vaccination (8). Systems vaccinology studies previously utilized microarray technology to identify transcriptomes. However, microarrays have now been virtually replaced by RNA-Seq technology due to its technical superiority (larger dynamic range, no signal saturation, and no restriction to a static set of printed probes) (9-12). Unlike microarrays, RNA-Seq is inherently flexible, and several technical parameters can be tailored uniquely for each experiment. The goal of this study was to establish parameters to optimally apply RNA-Seq to clinical vaccine studies.

Some work has been done to benchmark RNA-Seq analysis and understand how technical parameters influence experimental findings (13-18). Quality metrics, methods to estimate reproducibility, and algorithms to estimate statistical power have been developed for RNA-Seq technology; however, these have almost exclusively been defined using standardized reference samples (14, 19-22). Despite these efforts, relatively little work has been done to benchmark the utility of reference controls (Universal Human Reference RNA (UHRRs)), External Reference Control Consortium (ERCC) synthetic RNA spike-ins) and to assess the impact of the choice of coverage and read length in the context of “real-life” biological samples, or RNA-Seq-based clinical immunology/vaccine studies (23-26). In this study, we sought to empirically establish the impact of these and other technical parameters on the ability to accurately assess differential expressed genes (DEGs) using samples collected for a phase II clinical trial of a live attenuated tularemia vaccine. While we provide a high-level summary of transcriptomic changes following vaccination, this study was not intended to be a comprehensive biological transcriptional characterization of Tularemia vaccination or to identify correlates of Tularemia vaccine protection; these studies recently have been published elsewhere (27, 28). Specifically, our main goals were to: (i) assess the reproducibility of gene expression measurements and the ability to detect the same DEGs in two different laboratories using technical replicate samples; (ii) assess the impact of sequencing depth on gene representation of the global PBMC transcriptome using pooled, ultra-deep transcriptomes; (iii) estimate statistical power to determine DEGs as a function of effect size, sequencing coverage, and sample size; (iv) define the impact of changes in technical parameters (coverage depth and read length) on identification of DEGs; (v) assess the accuracy of fold-change estimates using synthetic mRNA spike-ins for varying read-count filterings cutoffs; (vi) determine an empirical read-count cutoff for filtering out lowly-expressed genes; and (vii) establish recommendations for RNA-Seq analyses conducted in clinical vaccine studies. It is also important to note that the goals of this study were not to perform an exhaustive comparison of established analytical methods to quantify gene expression or detect DEGs, or to compare the performance of these tools, as these comparisons already exist in the literature (29). The replication of the study at different sites was performed in a manner similar to that described in the influential “SEQC/MAQ-III” studies that sequenced a small number of technical replicates at multiple sites (16). The rationale for replicating this benchmarking experiment on the same sample sets (parallel aliquots of PBMCs from the same subjects and time points) run in two separate laboratories was to firmly establish if the findings could be validated using “real world” clinical trial samples.

## Results

### Experimental Design

The RNA-Seq benchmarking study described here utilized samples from a Phase 2, multi-center, double-blind randomized trial comparing the immunogenicity of two live attenuated vaccines against *Francisella tularensis*: an aging stock used for decades by the United States Army Medical Research Institute of Infectious Diseases (USAMRIID-LVS) and a novel lot produced by the Dynport Vaccine Company, DVC-LVS, intended to be its replacement (30) (**Figure 1)** (registered at ClinicalTrials.gov NCT01150695) (7). Replicate aliquots of the same PBMC sample from each subject time point were used to evaluate the agreement between laboratory RNA-Seq results (**Figure 1B**). Briefly, RNA was extracted from PBMCs from 10 healthy subjects at Days 0 (pre-vaccination), 1, 2, 7, and 14 following DVC-LVS vaccination at both sites (referred to as Site 1 and Site 2). Samples were prepared for sequencing using poly A selection followed by mRNA fragmentation, reverse transcription, adapter ligation and amplification at two different sequencing facilities; Site 1 employed Illumina TruSeq and Site 2 utilized a similar “in house” protocol(see Materials & Methods). A single operator at each site extracted RNA, conducted the library preparation, and performed CBot clustering and loading of pooled libraries onto Illumina HiSeq 3000 sequencers. Internal (ERCC) and external (UHRR) RNA controls were included in each experiment to estimate the dynamic range and fold change accuracy, as well as to evaluate the suitability of external RNA references for inter-site normalization. RNA sequencing was performed at each site by distributing the libraries for 53 samples evenly across 5 lanes of the respective Illumina HiSeq 3000 devices at each site (**Figure 1B**). Sequences were aligned to the GRCh38 reference genome.

**Figure 1:**
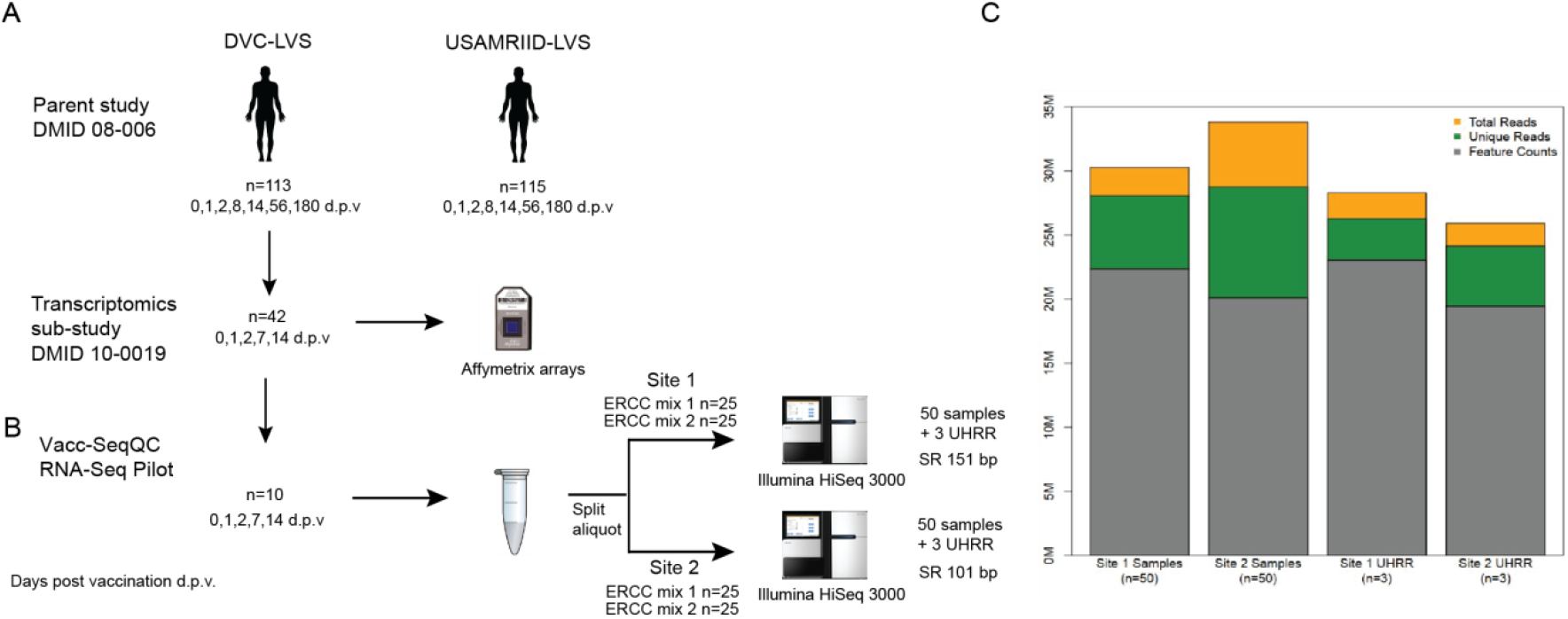
Experimental design of RNA-Seq benchmarking for the clinical vaccine study. **(A)** The samples utilized in this benchmarking experiment were obtained from parent study DMID 08-006 (ClinicalTrials.gov NCT01150695), of which 42 patients had RNA samples collected for microarray analysis. (**B**) RNA from five time points taken from 10 volunteers, for a total of 50 samples, at both Site 1 and Site 2. Each biological sample was spiked with either ERCC mix 1 or mix 2, for 25 replicate spike-ins of each mix. Samples were sequenced as single-ended 151 or 100 bp reads on an Illumina HiSeq3000 at each Site, with multiplexing targeting a depth of 30 M reads. At each site, an additional set of three RNA samples from UHRR controls were included for an independent reference. (**C**) Median sequencing metrics for all samples at each site. The yellow bar indicates the median total read depth obtained for all samples for each site; the top of the green bar indicates the total reads that are uniquely mapping to the GRCh38 reference, and the grey bar indicates the reads mapping to annotated features (i.e., genes) in the reference.

### Sequencing Statistics

A summary of the reference alignment statistics is listed in **Table 1**. Site 1 data was sequenced at 151 nt singled ended (SE) to a median total depth of 31.8 × 10^6^ reads, with 28.1 × 10^6^ (93.8%) uniquely mapping reads, whereas Site 2 data was sequenced at 100 nt SE and had a median total and unique read depth of 34.9 × 10^6^ and 28.8 × 10^6^ (88.1%), respectively (**Figure 1C and Table 1**). When reads were mapped against known gene models, a median of 22.4 × 10^6^ and 20.1 × 10^6^ reads were uniquely counted in the expression quantification step for Site 1 and Site 2 data, respectively. The majority (median of 86.6% for Site 1 and 76.6% for Site 2) of tags (i.e. spliced reads) mapped to known exonic regions, and fewer mapped to intronic (12.6% and 22.2%, respectively), and intergenic regions (0.9% and 1.2%, respectively) (**Table 1**). The GC content in sequenced libraries was comparable between sites (medians 49.3, 50.5 for Site 1 and Site 2 data, respectively). Of note, one baseline (pre-vaccination) sample at Site 1 showed substantial GC content bias with a median GC of 55.6% and was identified as an outlier in principal component analysis (PCA). This sample was removed from downstream analyses unless otherwise specified.

**Table 1:** Summary of human reference alignment statistics.

| Sequencing Parameter* | Site 1 (N=50) |  |  |  |  | Site 2 (N=50) |  |  |  |  |
| --- | --- | --- | --- | --- | --- | --- | --- | --- | --- | --- |
|  | Min | Q1 | Median | Q3 | Max | Min | Q1 | Median | Q3 | Max |
| Total Reads [$10^6$] | 26.69 | 30.39 | 31.81 | 34.25 | 42.69 | 21.16 | 30.32 | 34.85 | 43.95 | 63.76 |
| Total Mapped Reads [$10^6$] | 25.37 | 28.71 | 30.27 | 32.54 | 40.65 | 20.34 | 29.27 | 33.8 | 42.43 | 61.65 |
| Unmapped Reads [$10^6$] | 1.0 | 1.5 | 1.61 | 1.71 | 2.05 | 0.66 | 1.05 | 1.19 | 1.49 | 2.11 |
| Uniquely Mapped Reads [$10^6$] | 16.06 | 26.69 | 28.06 | 30.19 | 37.47 | 18.22 | 25.92 | 28.76 | 36.79 | 55.74 |
| Uniquely Mapped Reads [%] | 42.7** | 92.8 | 93.75 | 94.2 | 95.3 | 83.7 | 86.3 | 88.1 | 90 | 91.1 |
| Counted Reads [$10^6$]*** | 11.65 | 21.51 | 22.36 | 24.35 | 30.24 | 12.32 | 17.57 | 20.1 | 25.16 | 37.13 |
| Median GC [%] | 47.83 | 49.01 | 49.33 | 49.67 | 55.63 | 49.5 | 50.5 | 50.5 | 50.5 | 51.49 |
| Exon Tags [%] | 64.33 | 85.57 | 86.6 | 87.42 | 91.44 | 71.95 | 75 | 76.61 | 77.99 | 82.63 |
| Intron Tags [%] | 8.04 | 11.64 | 12.6 | 13.55 | 33.45 | 16.38 | 20.9 | 22.16 | 23.74 | 26.82 |
| Intergenic Tags [%] | 0.52 | 0.79 | 0.85 | 0.89 | 2.21 | 0.99 | 1.11 | 1.2 | 1.25 | 1.34 |

Taken together, these summary statistics demonstrated that, in general, the replicate samples were sequenced effectively at each site. However, despite using replicate PBMC samples as starting material and identical sequencing platforms, coverage metrics and mapping statistics varied. Variation was less pronounced for the 3 external UHRR control samples whose sequences were combined prior to processing (**Figure 1**) indicating that variability in RNA content between paired samples may have contributed to this. In addition, read lengths differed between protocols (151 vs. 101 nt). Except for read length, we considered these differences between datasets to reflect true variability introduced by different operators generating distinct library preparations. Thus, these data should represent a good comparison set for investigating the overall reproducibility of Illumina-based RNA-Seq results in the context of a vaccine study.

### Inter-site agreement of RNA-Seq log_2_ counts per million and fold-change estimates improved after filtering out lowly-expressed genes

A prior study assessed the technical accuracy of RNA-Seq measurements between laboratory sites (14). This prior study relied on comparison of read counts of reference RNAs (i.e. UHRR) between laboratories, which involved non-matching sequencing platforms. Here, we extended this benchmarking of RNA-Seq accuracy by using replicate samples from clinical samples. To assess the agreement of read-count estimation and fold change, and to test the impact of filtering lowly-expressed genes, we calculated an adjusted Euclidean distance (which uses the mean squared distance rather than the sum of squared distances to normalize for the total number of genes) and Pearson correlation between the two sites for the log_2_ counts per million (LCPM) (n=49) and the log_2_ fold change (LFC) (n=36, post-vaccination vs. pre-vaccination) for each subject. This analysis was performed on the data after applying different counts-per-million (CPM) filtering cutoffs (unfiltered, 1, 2, 4, and 8 CPM) to remove lowly-expressed genes; these filtering cutoffs were applied by removing any gene whose maximum expression level across samples was below the specified threshold.

The Euclidean distance showed improved reproducibility of gene expression profiles between replicate samples as the CPM cutoff increased (**Figure 2A)**; the average Euclidean distance was 1.45 for the unfiltered dataset, and this distance dropped to 0.80 after applying the >1 CPM cutoff. Increasing the threshold to >2 CPM, >4 CPM, and >8 CPM showed additional, incremental shifts in the distribution toward lower Euclidean distances. Similarly, the Pearson correlation showed improved reproducibility with filtering; the mean correlation increased from 0.945 for the unfiltered data up to 0.960 for the >1 CPM filtering, but it did not continue to increase substantially with additional filtering (**Figure 2B**). The Euclidean distances of the LFC profile for each replicate paired sample showed a similar trend: reproducibility between Sites improved with increased filtering (**Figure 2C**). The average Euclidean distances on the LFCs shrunk from 1.06 for the unfiltered dataset down to 0.30 for the >8 CPM filtering. In contrast to the LCPM profiles, the Pearson correlation of the LFCs did show continued improvement with additional filtering (**Figure 2D**). The average Pearson correlation jumped from 0.11 for the unfiltered data to 0.48 with >1 CPM filtering. Increasing the threshold continued to improve the correlation reaching an average of 0.73 with >8 CPM filtering. A similar analysis was performed looking at reproducibility by gene (rather than by sample), and this also demonstrated improvements in Euclidean distance and Pearson correlation after applying filtering (**Supplemental Figure 1**).Taken together, these results demonstrated that (i) the overall concordance of log_2_ counts per million and fold-change estimates increased with increasing minimum gene expression abundance, (ii) low expressed genes provide inaccurate measurements, as demonstrated here by low concordance of data between sites, and (iii) commonly used cutoffs such as >1 CPM might not be sufficiently stringent to ensure accurate calculation of fold changes.

**Figure 2:**
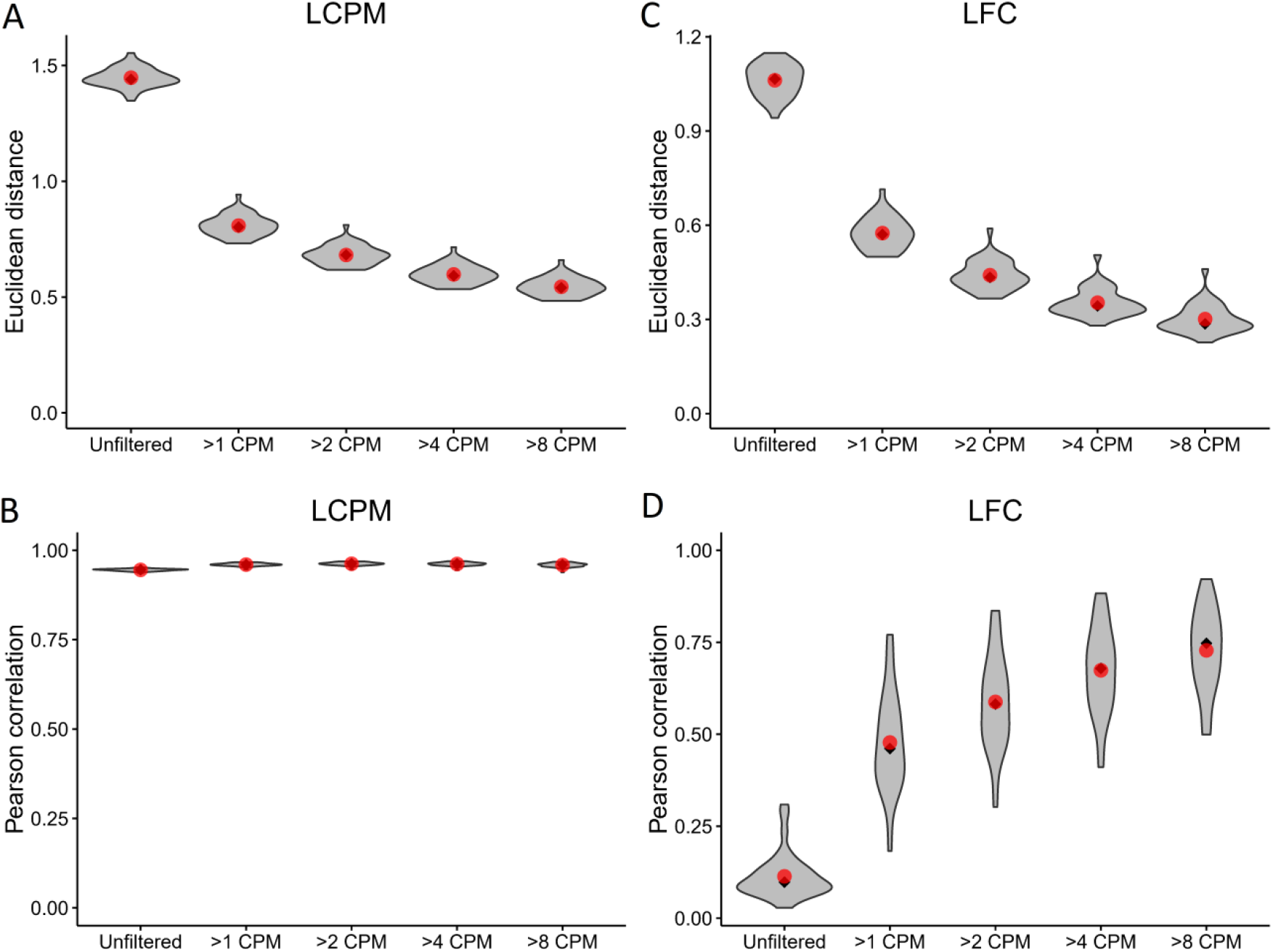
Inter-site agreement of RNA-Seq log_2_ counts per million and fold-change estimates improves after filtering out lowly-expressed genes. LCPM: log_2_ counts per million. LFC: log_2_ fold change. **(A)** Violin plot depicting the distribution of adjusted Euclidean distance between Site 1 and Site 2 of the LCPM gene expression profile for each subject. Each distribution consists of a total of 49 observations. Red dots mark the mean of the distribution, and black diamonds mark the median. Each column shows the distribution after filtering genes at the CPM thresholds indicated on the x-axis. **(B)** The same analysis and plotting design as described in (A), but for the Pearson correlation between Site 1 and Site 2 of the LCPM gene expression profile for each subject. **(C)** and **(D)** show the same results but calculated on the LFCs. For these, each distribution consists of a total of 36 observations.

### Discordance of detection of DE genes between sites was mainly driven by differences in mean fold change

To assess and compare the strength of the biological signal, i.e. the change in magnitude of gene expression longitudinally over time (Days 1, 2, 7 and 14), we contrasted mean LFC and mean LCPM using MA plots (**Figure 3A**). We determined the number of DEGs using edgeR (FDR <0.05, fold change of ±1.5) after applying a maximum CPM ≤8 across samples to filter out lowly-expressed genes. For both sites, an increase in fold change and the number of DEGs was observed over time indicating that the complexity of vaccine-induced gene expression signals in PBMCs reached a peak at Day 14. As shown in **Figure 3A**, substantially higher numbers of DEGs were detected at Day 14 as well as Day 7, compared to Day 1 and Day 2 for which few DEGs were detected. These results are consistent with other studies describing use of transcriptomics to characterize responses to live-attenuated vaccines, in which increases in gene expression corresponds with the initiation of the adaptive immune response. Overall, 1173 and 1218 DEGs were detected across all time-points for Site 1 and Site 2, respectively, of which 869 overlapped with a concordance of 74% for Site 1 and 71% for Site 2. To better understand where the loss of concordance in DEGs occurred, we examined whether missed DEGs were due to missing FDR and/or fold-change cutoffs or due to CPM filtering (**Figure 3A, Supplemental Table 1**). At the Day 1 and Day 2 time points, the overall DEG concordance was only ∼60% (Figure 3A, right panel, green bar). However, when we accounted for genes that had a significant FDR but did not pass the fold-change threshold of ±1.5, the maximum concordance increased to nearly 80% (**Figure 3A**, right panel, grey bar). Generally, we noted that the failure of genes to pass a FC cutoff (grey bars) was consistently the largest contributor to gene list discordance, followed by the effects of different levels of CPM filtering (blue bars) (**Figure 3A)**. However, failing both FDR and FC cutoffs had more pronounced effects than CPM filtering for the weaker biological signals observed at Days 1 and 2, indicating that statistical power was more negatively impacted by lower effect sizes and higher variability in changes for these days. Overall, only a small fraction of genes was discordant due to failing to pass only the FDR cutoff (orange bars). We considered that the lack of overlap due to fold change differences between Site 1 and Site 2 was due to the low magnitude of fold change for DEGs at Day 1 and Day 2. However, when we compared the Site 1 vs. Site 2 DEG overlaps from the Day 7 and Day 14 timepoints, which had more gene expression changes, the fraction of discordant genes due to failing to meet the fold-change criteria (grey bars) was similar. To examine the disagreement between fold changes in more in depth, we ranked the DEGs by fold change, from most downregulated to most upregulated, and plotted them for each post-vaccination day (**Figure 3B**). DEG fold-change ranks were highly concordant between the two sites and overall directions agreed. Variability in ranks was greatest for the middle ranks and strongest discordancy in ranks was mainly observed for genes at one site that had a fold change below the threshold of ±1.5 relative to the other site. These data were instructive for establishing the relative contribution of cutoffs to the discordance of DEGs between Sites. We demonstrated here that utilizing a lower FC cutoff of 1.5 can result in a reduction in inter-site agreement of DEGs by 20 percentage points compared with using higher FC cutoffs.

**Figure 3A:**
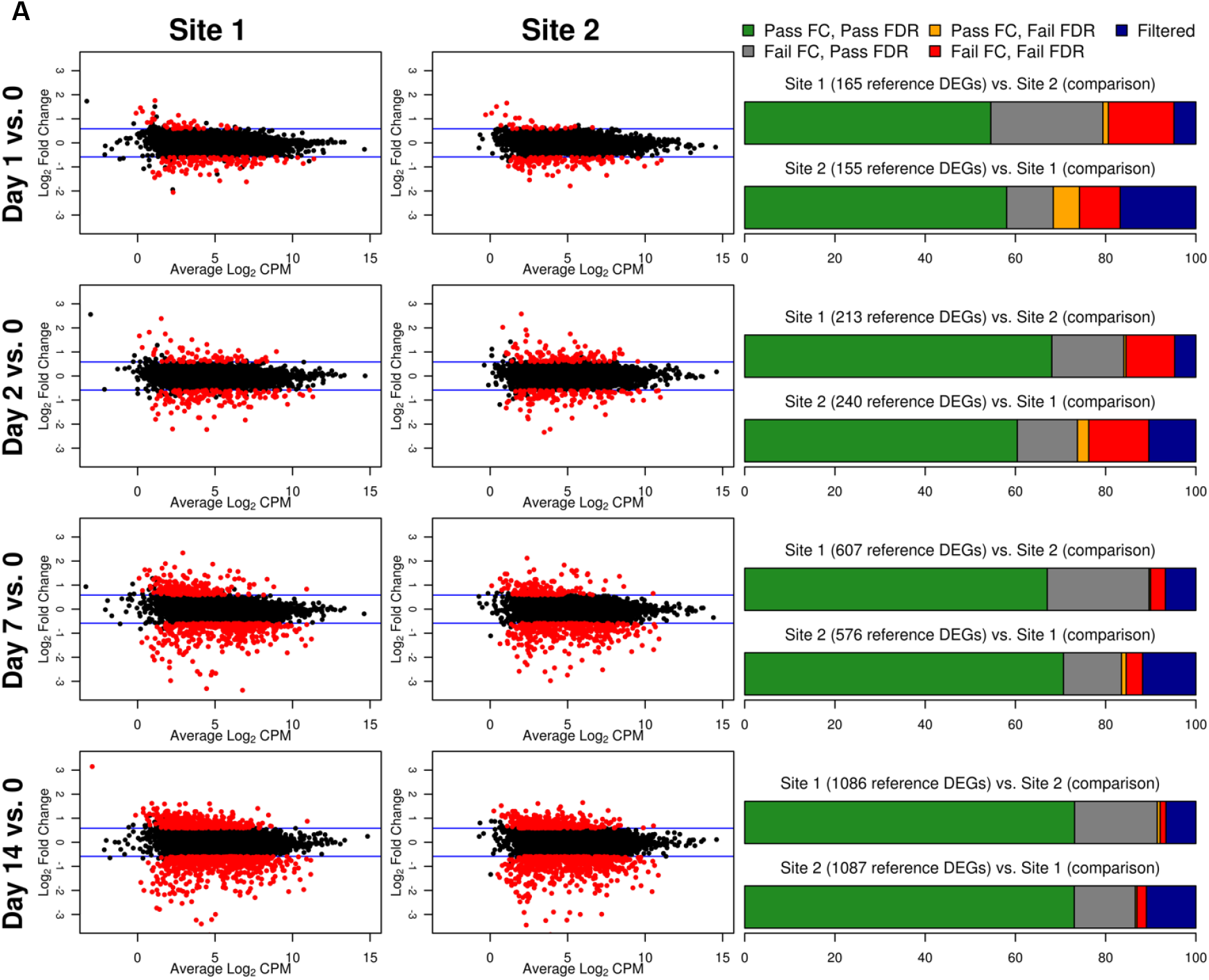
Fold change and DEGs over time. MA plots shown to left contrast average log_2_ fold changes from pre-vaccination (y-axis) by average gene expression levels (log_2_ CPM as shown on the x-axis) for each site and post-vaccination time point. Blue lines indicate the pre-specified minimum fold-change cutoff of ±1.5 fold. DEGs (edgeR FDR <0.05, fold change ≥ ±1.5 in either direction, and maximum read count of >8 CPM across all samples) are colored in red. Stacked bar plots to the right summarize the overlap between the two sites. The first bar plot for each post-vaccination day represents DEGs identified for Site 1 color-coded by the overlap class for genes identified for Site 2 (combination of FDR and fold change criteria or CPM gene filtering criterion that were met/not met). The second barplot shows the reverse presenting a characterization of Site 2 DEGs based on the overlap with Site 1 genes.

**Figure 3B:**
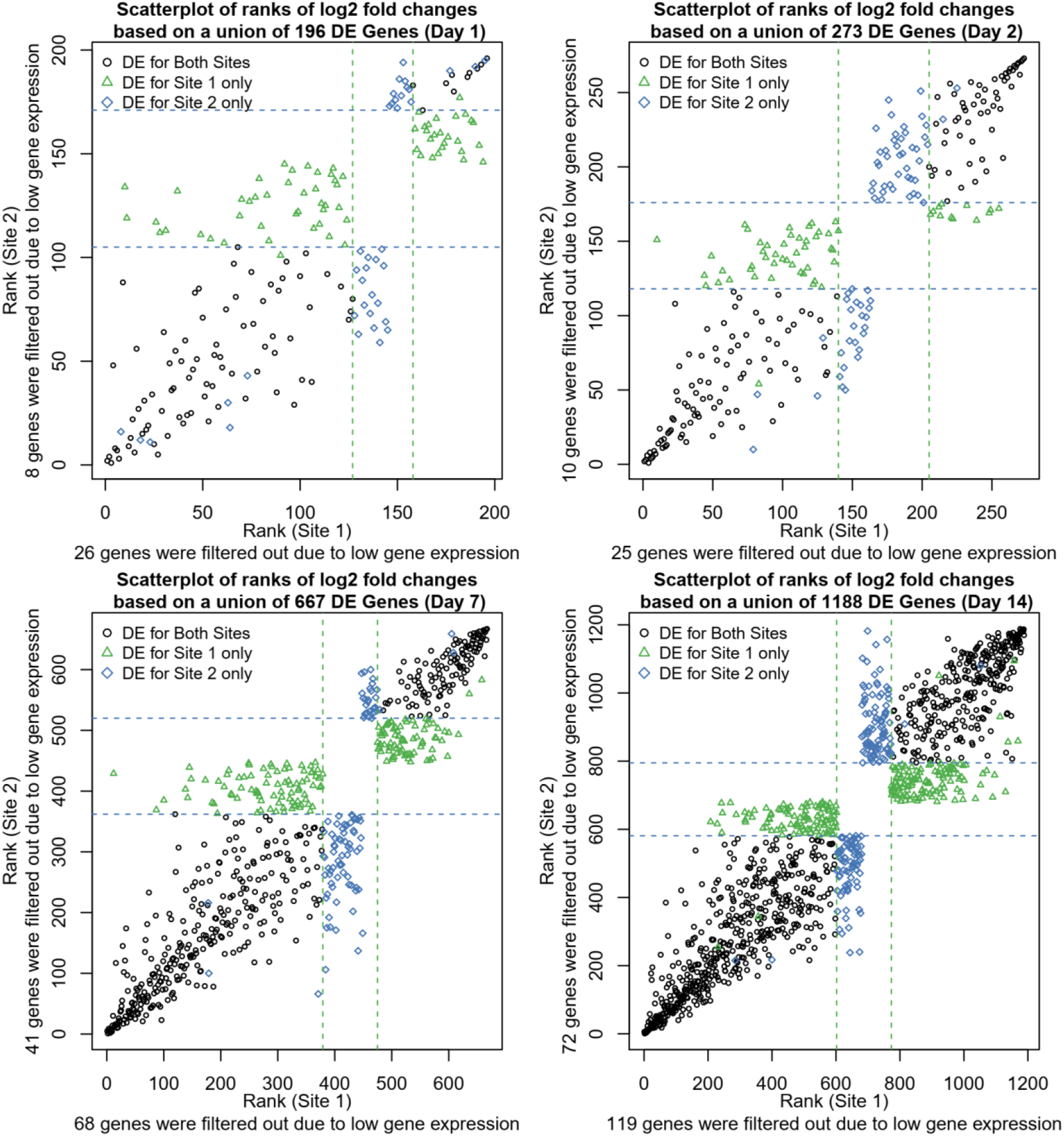
Scatterplots of ranks of log_2_ fold changes based on a union of DEGs identified for each site by post-vaccination day. A rank of 1 corresponds to the lowest log_2_ fold change. Dashed lines represent the upper and lower rank boundaries of the ≥1.5 fold-change cutoff for each site.

### The influence of sequencing depth on gene representation of the PBMC transcriptome

We next assessed the influence of sequencing coverage (read depth) on gene representation in our datasets. To establish a comprehensive reference of genes expressed in PBMCs, we first pooled, *in silico*, data from all 100 samples (10 individuals, 5 time points, 2 sequencing sites), yielding a reference transcriptome comprising 2.2 B total reads that mapped to hg38 human reference gene models. We next filtered out genes with fewer than 100 total aggregate reads (corresponding to approximately 1 read per sample). This resulted in a “PBMC transcriptome compendium” comprised of 26,172 gene models that we used as a reference set considered “truly expressed” in the PBMC compartment. To assess the impact of coverage on gene representation, we used the PBMC transcriptome compendium to determine the proportion of genes detectable per sample as a function of decreasing simulated coverage (i.e., read depth) (**Figure 4**). To focus on genes with strong evidence of genuine expression, we choose a cutoff of ≥16 reads per sample. At a depth of 30 M reads, an average of 14,545 genes were detected with ≥16 reads, representing 56% of the truly expressed PBMC compartment (or 44% lost). While relaxing our read-count threshold increased the number of detected genes, there was still incomplete representation of transcripts: at a threshold of 5 reads, only 67% of genes in the compendium were detected, and at a threshold of 1 read, detection was 83%. As expected, the number of detected genes from the PBMC reference transcriptome compendium decreased with lower coverage. At 25 M reads, the fraction of genes with ≥16 reads from the cumulative PBMC reference was 53%, and at 20 M reads the fraction of genes lost was 52%. In comparison, when subsampling *in silico* to 10 M reads per sample, 12,030 genes (46%) were detected on average, representing a loss of 54% relative to the cumulative gene set. At coverage of 10 M reads, on average 2,515 genes (9.6%) fewer genes with ≥16 reads could be detected compared to samples at 30 M. This demonstrated that, even at relatively deep sequencing coverage of 30 M, a significant percentage of the gene content expressed in PBMCs was not detected.

**Figure 4:**
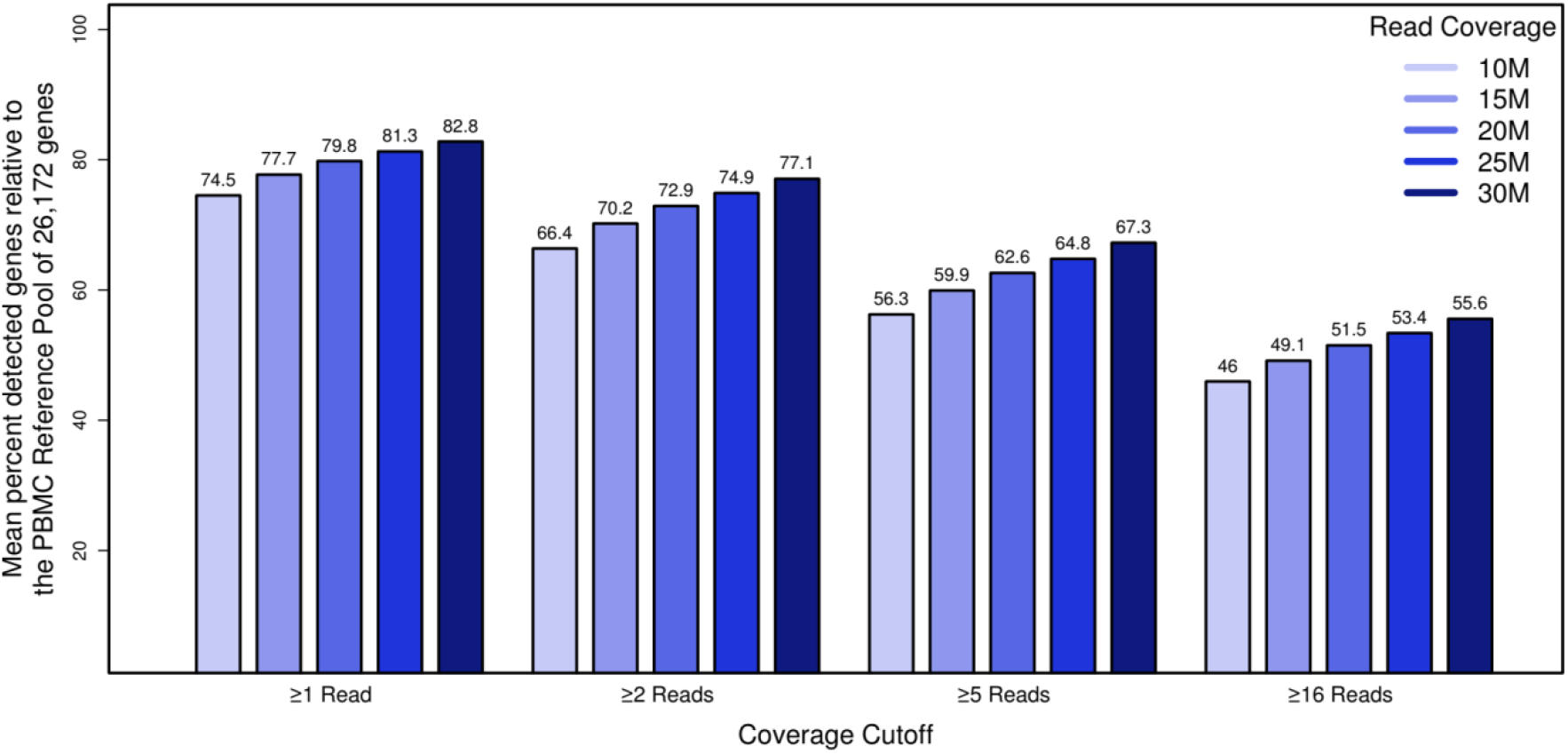
The impact of read depth on gene representation by establishment of a deep “compendium” transcriptome of PBMCs. The x-axis represents simulated datasets (n=50) with coverage x, the y-axis shows the percentage of genes that we considered truly expressed in the PBMC compartment for genes that met the filtering cutoff. Bar plots summarizing the mean percentage of the detected PBMC compartment genes by coverage and read filter criterion.

### Coverage was more influential than read length in ensuring consistent DEG results

Two of the main technical parameters that determine cost in an RNA-Seq experiment are (i) read-length and (ii) sequencing depth. The impact of read length on the ability to accurately measure DEGs has not been comprehensively studied to date: One previous study utilized data from the SEQC study and determined virtually no impact of using paired end reads on the ability to reproducibly detect DEGs defined by p-value (17). Thus, we focused our analysis on defining the optimal single-ended read length. In the Chhangawala study (17), the dataset utilized commercial reference RNA samples (UHRR and Ambion FirstChoice Human Brain Reference RNA) and relatively low replication (n=3). In this regard, the current study, which used “real-world” samples from a vaccine trial and was more robustly powered (n=10), provided an optimal opportunity to empirically test the impact of varying read-length parameter. To evaluate the impact of different read lengths on DEG identification, we used the Site 1 dataset as it utilized longer read lengths compared to Site 2. Different read lengths were simulated by right-truncating the actual sequencing reads (100 nt, 75 nt, and 50 nt read length) to most accurately mimic true Illumina sequencing and the range of Q-scores that would result from bona-fide sequencing at those lengths. To simulate different coverage levels, we randomly down-sampled the original 31.8 M reads to 25 M, 20 M, 15 M, and 10 M reads. For the comparisons, we used the DEG results identified for the original data (151 nt read length and 31.8 M read coverage) as the reference, essentially controlling for all other parameters. We then compared DEGs obtained using differing read lengths and coverage levels using the Jaccard index. Results showed that shortening read length had a lesser impact on DEG detection compared to reduction in sequencing depth (**Figure 5**). Notably, although the set of genes identified as DE was affected, the total number of DEGs detected did not change substantially with either shortening read length or a reduction in sequencing depth (**Supplemental Figure 2**). Reducing the read-length from 151 to 100, 75, and 50 nt at the same coverage level (35 million) showed a mean DE agreement (based on Jaccard index) for days with strongest biological signals (Day 7-14) of 0.95, 0.94, and 0.91, respectively. At the lowest read length level of 50 nt, 91% of the detected DEGs agreed. In contrast, reducing coverage from 35 to 25, 20, 15, and 10 M reads at the same read length (151 nt) showed a mean DE agreement (based on Jaccard index) for days with strongest biological signals (Day 7 and 14) of 0.93, 0.87, 0.83, and 0.80, respectively (**Figure 5**). In summary, this showed that read length had a much lesser impact on DEG agreement compared to coverage. Changing the read length by approximately half from 151 nt to 75 nt only resulted in 6% disagreeing DEGs. In contrast, reducing coverage by approximately half from 32 M to 15 M reads resulted in 17% disagreeing DEGs. The disagreement was not explained by a change in the total number of DEGs.

**Figure 5:**
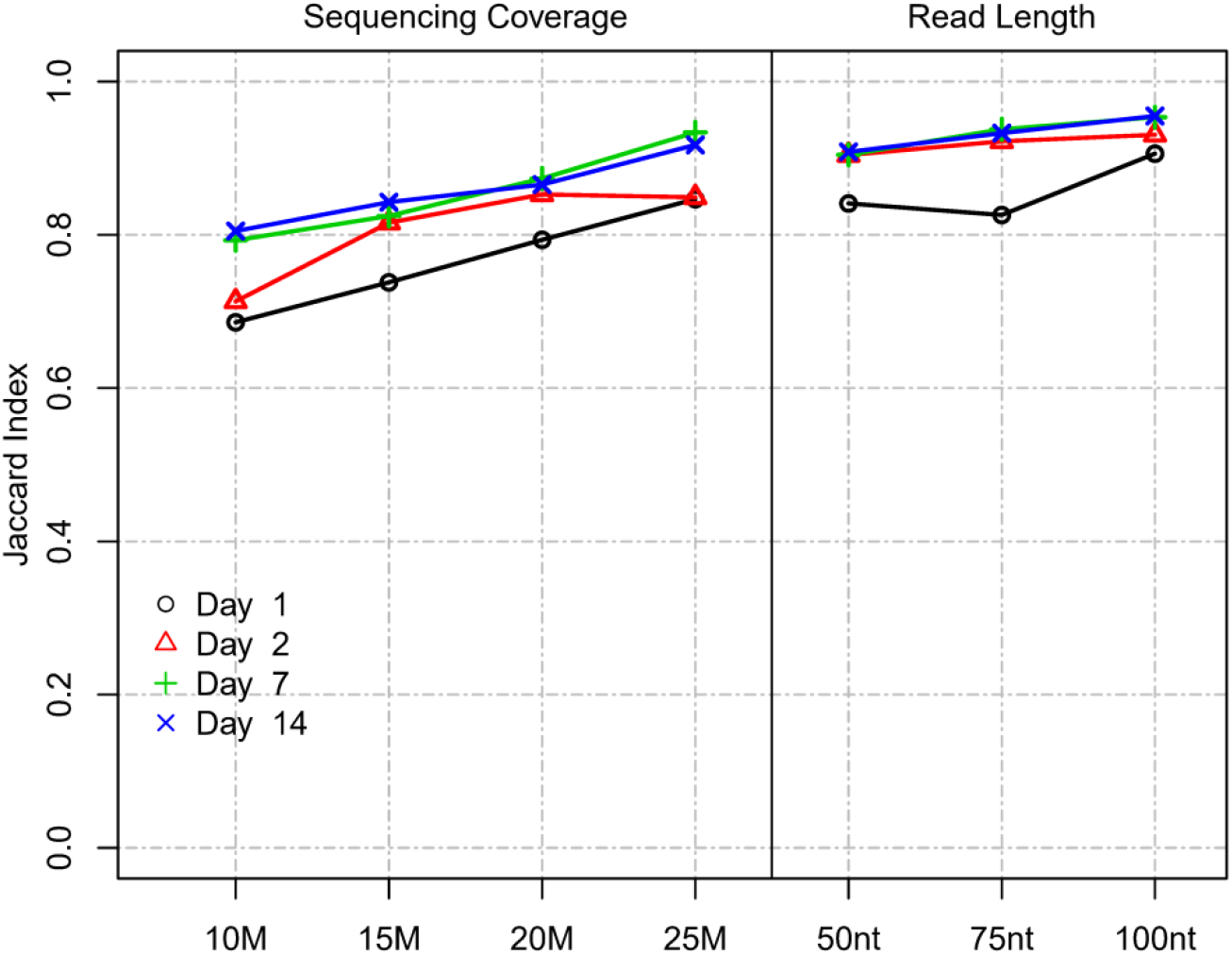
Impact of read length and coverage on DEG detection. Impact of read length and coverage on Jaccard index between DEGs (Site 1). Jaccard index represents the proportion of intersecting DEGs compared to the union of DEGs identified for a subsampled/truncated dataset and the original dataset.

### Establishment of a read-count cutoff for accurate fold-change assessment

In RNA-Seq experiments, pre-filtering genes with low read counts to reduce low quality data is an important part of the analysis but is frequently done using arbitrarily defined thresholds. As we have shown in **Figure 2**, filtering also improved inter-site agreement in LCPM and LFC based on Pearson correlation. Exogenous RNA species used as spike-in controls provide the ability to include within a dataset a set of references with known abundance levels that can be used as an internal “truth” by which to judge the accuracy of the analysis accounting for varying sequencing parameters. Here, we tested the utility of the spike-in controls provided by the External RNA Controls Consortium (ERCC) (24) to empirically derive read-count cutoffs within our datasets that result in a high agreement between expected and observed fold changes. To achieve this, we used 92 ERCC spike-in RNA control pairs (ERCC2 vs. ERCC1 mixes with known abundance ratios) that were spiked into pre-defined sample pairs (n=40 for each site). The 92 spike-in transcripts were binned into 7 groups of increasing expression levels (based on the average LCPM of each ERCC1 and ERCC2 spike-in pair), and the Spearman correlation between the observed and expected ERCC 2 vs. ERCC 1 ratios was calculated within each bin (on average 13 ERCC1/2 spike ins per bin) for each paired sample, along with the respective average LCPM. These values were used to model the relationship between the LCPM and the correlation of empirical vs. expected fold changes (**Figure 6**). The trend plots showed that the correlation increased with increasing LCPM values. We then empirically determined a read-count threshold based on LCPM abundance that marks good agreement between expected and observed LFC. We defined good agreement as having a correlation value of 0.9 or higher. The non-linear relationship between Spearman correlation and average LCPM was modeled using a 3^rd^ order polynomial. The model was fit to our two datasets (n=40 paired samples each), and the LCPM value that gave a predicted correlation value of 0.90 was used as the empirically determined threshold. This method yielded LCPM expression cutoffs of 3.34 and 1.73 for Sites 1 and 2, respectively. To assess the degree of variability of the point estimate we used bootstrapping. Resulting 95% CIs were 2.15 to 6.84 LCPM forSite 1 and 1.29 to 2.17 LCPM for Site 2 (**Figure 6**). These intervals can be used to balance the trade-off between fold change accuracy and filtering out too many genes. To maximize the number of genes for the analysis while maintaining good agreement between observed and expected fold changes, the lower bound of the 95% CI may be used as the cutoff threshold, in this case 2.15 and 1.29 for Site 1 and Site 2, respectively. With nearest integer increments typically used for read counts, this would translate to 3 and 2 LCPMs for Sites 1 and 2, respectively.

**Figure 6:**
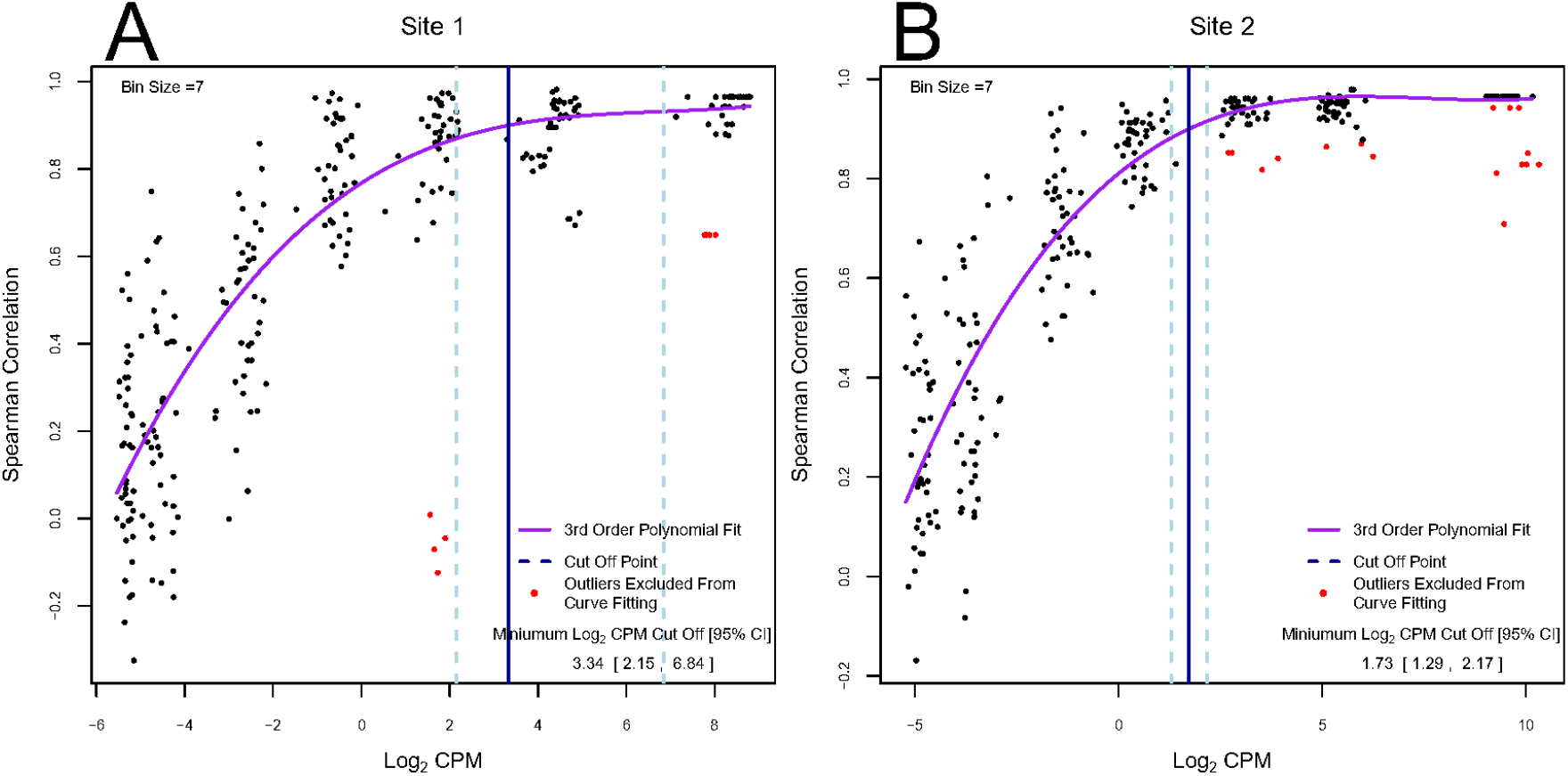
Determination of empirical minimum expression cutoffs using ERCC spike-ins. Each dot represents a subset of the 92 ERCC1 and ERCC2 transcripts for one paired sample grouped by average ERCC1 and ERCC2 abundance (on average 13 per abundance bin). The y-axis summarizes the Spearman correlation between the empirical log_2_ fold change and expected log_2_ fold change across these transcripts. The x-axis represents the average log_2_ abundance per paired sample for a subset of the ERCCs. Overall, 40 pre- and post-treatment pairs were created for each experiment. Each pair is shown multiple times (once in each abundance bin). The purple line represents the fitted 3^rd^ order polynomial function. Red dots represent paired samples with outlying correlation results relative to their abundance bin (exceeding ±1.5 times the interquartile range) which were excluded from curve fitting. The vertical blue solid line indicates the log_2_ counts per million value at which the fitted curve is equal to a correlation value of 0.9, and the vertical light-blue dashed lines show the 95% bootstrap confidence interval for this log_2_ counts per million value. The values in the lower right correspond to these vertical lines: the point estimate and 95% bootstrap confidence interval of the log_2_ counts per million at which the fitted polynomial was equal to 0.9.

In this section we highlighted a new algorithm that utilizes internal ERCC spike-ins for determining an optimal read-count threshold for each dataset independently. The advantage of this approach is that it is data driven and implicitly controls for any inherent sequencing parameters. These results suggested that the 8 CPM cutoff (3 LCPM) used in the original analysis of these data provided a good agreement between observed and expected fold changes for Site 1. For Site 2, a 4 CPM cutoff (2 LCPM) would have been more optimal.

### Power estimates for vaccine studies for DEG detection

To further evaluate the impact of different parameters on DEG identification, we assessed relative statistical power for different scenarios characterized by varying sample size (n=3 to n=15), effect size (1.25 to 2 minimum absolute fold change), and read coverage (10 to 60 M reads) (**Figure 7**). In lieu of a “truth set” of genes for which the differential expression was independently known, we defined a set of “truly differentially expressed genes” (TDEGs) as those genes identified as DEG for both sites using two different R packages (edgeR and DESeq2). Using these conservative criteria, we obtained 444, 357, 2,300, and 3,284 TDEGs for post-vaccination Days 1, 2, 7 and 14 respectively. We then simulated negative binomial data for each post-vaccination day using Site 2 data as the basis for power analyses as this set was more complete than Site 1 due to one outlying subject.

**Figure 7:**
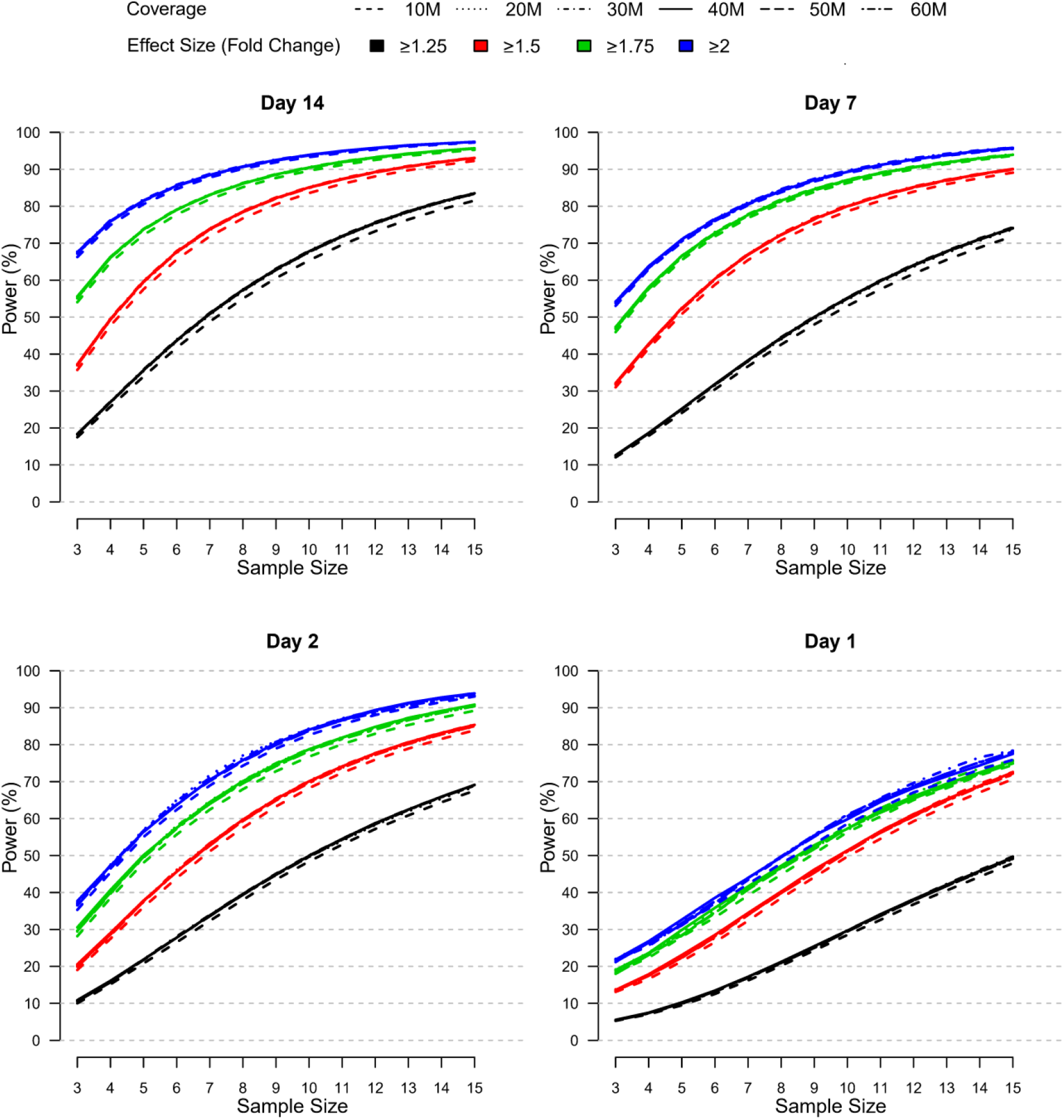
Relative power by samples size, effect size, and coverage at each post-vaccination day as simulated using the modified PROPER R package. Days were sorted by decreasing vaccination effect based on overall fold changes and DEG responses observed for this study (see **Figure 3A**). Power was assessed for different fold-change cutoffs (indicated by color-coded lines), coverage (as indicated by the line type), and sample size (x-axis).

The power curves shown in **Figure 7** demonstrate that statistical power to detect TDEGs was most strongly influenced by sample size. Importantly, power was modulated by the overall magnitude and breadth of the biological vaccine effect, which was evident by the overall horizontal shifts in the power curves with increasing power for increasing vaccine effects reported for Days 1-14 (see **Figure 3A**). Most positively impacted was the statistical power to detect TDEGs with smaller fold changes (FC ≥1.25 fold change), for example, with a sample size of n=10 and coverage of 30 M reads, the statistical power increased from ∼30% on Day 1 to 50%, 55% and 68% on Days 2, 7, and 14, respectively. Three other notable trends were apparent:

First, statistical power to detect TDEGs was remarkably higher for genes with larger effect sizes. For example, at Day 14, for genes with greater than 1.75-fold or 2-fold effect sizes, even at the lowest level of sample size evaluated (n=3) the statistical power was 55% and 68%, respectively. An increase in sample size to 10 or higher led to greater than 90% power for detection of TDEGs with fold changes at or above 1.75 (**Figure 7**). In contrast, at a lower effect-size cutoff of 1.5-fold, the approximated power was 37% at a sample size of n=3, and 90% power wasn’t attained until a sample size of 13 or more. For the smallest effect size considered, a cutoff of 1.25-fold, statistical power dropped much lower for smaller n with only 18% of TDEGs detected at n=3. However, sensitivity increased steadily with increased sample sizes: at n=6, 44% of TDEGs were detected; at n=10, 68% of TDEGs were detected; and at n=15, the largest sample size considered, 84% of TDEGs were detected.

Second, read coverage had a minor effect on power (**Figure 7**). A noticeable negative effect on power was only observed when coverage was reduced from 20 M to 10 M, and even with this magnitude of coverage reduction, the power difference was minor. No substantial gains in statistical power were detected beyond 20 M reads. Furthermore, this lack of dependence of power on the read coverage depth was observed across all four days. This means, since the biological signal increased across time, deeper coverage over 20 M reads did not improve power in the presence of weaker biological signals.

Lastly, the false discovery rate (FDR), defined by the proportion of statistically significant genes that were not TDEGs, showed a strong dependence on the strength of the biological signal (**Supplemental Figure 3**). The statistical significance of each gene was determined based on an adjusted p-value set to control the false discovery rate at a 5% level. At Day 1, when the signal was weakest, the approximated FDR was inflated to 41% for a sample size of n=3. However, the FDR improved as the sample size increased; with a sample size of n=13 or larger, the FDR was below 10% for all fold-change cutoffs and below 5% for the more stringent 1.75- and 2-fold-change cutoffs. In contrast, at Day 14 when the biological signal was strongest, the FDR was near 5% for all sample sizes and fold-change cutoffs. Of note, the coverage level did not noticeably affect the FDR, regardless of the strength of the biological signal, similarly to what was observed for power.

The type-I-error rate, defined as the proportion of non-TDEGs that were declared statistically significant, was also evaluated (**Supplemental Figure 4**). In all simulation settings, the type-I-error rate was below 2%. It should be noted, however, that a low type-I-error rate is not surprising in this context because its value is directly related to the proportion of genes declared to be statistically significant. As a result, the type-I error should be assessed together with power to form an accurate summary of the performance. With this in mind, we found that: (i) the stronger biological signals at Days 7 and 14 led to increased type-I error relative to Days 1 and 2, but this was accompanied by improved power; (ii) there was no apparent dependence between coverage level and the type-I-error rate in any scenario; and (iii) the type-I-error rate remained relatively unchanged with increasing sample size, except at Days 7 and 14 when the lowest fold-change cutoff of 1.25 was used, in which case the type-I error showed a slight, gradual increase for each sample size considered even though power still improved.

While the magnitude of the fold change for the TDEGs was fixed in this simulation, the variance of these genes was allowed to change based on each simulated coverage dataset. Since variance plays an important role in power – higher variance directly results in lower power, all else equal – we investigated whether the mean-variance relationship across genes was affected by the different coverage depths. Overall, the mean-variance trends across genes were maintained for different coverage depths considered in this simulation; the curves shift left with lower coverage to produce fewer total counts, but the trend of lowly-expressed genes having higher variance compared to highly-expressed genes persisted (**Supplemental Figure 5A**). While the trends shifted vertically with increasing log mean expression, the corresponding log over dispersion values remained similar. Additionally, after applying TMM-normalization and obtaining LCPM values for the gene expression in the simulated coverage datasets, these values aligned with the LCPM values in the original dataset (**Supplemental Figure 5B**). Hence, this suggested that coverage depth would also have a minimal differential impact on lowly-expressed gene filtering based on CPM. Thus, while we performed this analysis for genes with > 8 CPM, given this result, our findings that coverage did not have a substantial impact on power can be generalized to other CPM cutoffs.

## Discussion

RNA-Seq offers a large amount of flexibility in the design of experiments, and many technical parameters such as read length, sequencing depth, and replication (i.e. sample size) can be varied to match the needs of an experiment. In addition, choices of lowly-expressed gene and fold-change thresholds have an impact on downstream analysis. The goal of most RNA-Seq experiments is to detect genes with statistically significant differences in expression across conditions. Several studies in recent years have sought to investigate how technical parameters influence the composition of DEGs and to provide standards for the design of RNA-Seq experiments. The contributions of technical variation to RNA-Seq analysis have been reduced in recent years due to the improvements in library preparation methods and widespread use of Illumina sequencing; however, the contribution of biological variation remains unique to each dataset.

With this in mind, we sought to provide general recommendations for conducting RNA-Seq in clinical vaccine studies using real-world data generated from a vaccine clinical trial. To achieve this, we assessed human blood PBMC transcriptomes for 10 healthy subjects at Days 0 (pre-vaccination), 1, 2, 7, and 14 following vaccinations with a *Francisella tularensis* live vaccine strain (DVC-LVS) and compared results obtained from matching aliquots from two different sequencing laboratories addressing the following questions:

1. ***Inter-site reproducibility:*** *How reproducible are data generated in separate laboratories? How can agreement be improved?*
2. ***Technical Parameters***: *What is the impact of altering lowly-expressed gene thresholds, read length, and coverage? How is PBMC transcript coverage impacted? How can lowly-expressed gene thresholds be derived empirically?*
3. ***Sample Size & Statistical Power***: *Given the biological variation and effect sizes observed in a clinical vaccine study, what relative role does sample size, effect size, and coverage play? What is the impact on false discoveries and the type 1 error?*

### Inter-site reproducibility

The seminal SEQC study investigated concordance of samples processed in multiple sites performing contrasts of two reference samples (Universal Human Reference RNA vs Brain Reference RNA). One important conclusion from SEQC was that while RNA-Seq fold-change estimates were generally concordant between laboratories, read-count data could not be reliably combined (16). Normalization improved overall inter-site agreement but not to the level where read counts could be combined (31).

In this study, we showed that the inter-site agreement between log_2_ count per million and fold changes was strongly dependent on the degree of read-count filtering. While the commonly used cutoff of >1 CPM resulted in the biggest gain, inter-site agreement showed continuous further improvements with more stringent filtering and observed Pearson correlation was modest with an average of r=0.7 for LCPM and r=0.50 between LFCs. This indicated that (i) genes that are lowly expressed disproportionally contribute to disagreement between sites, as demonstrated here by lower concordance of data between sites when not excluded; (ii) that these low abundance transcripts are most effectively negated by filtering; (iii) average explained variance was 92% and 24% for LCPM and LFC, respectively, when using >1 CPM demonstrating that a lot of the inter-site variability remained unexplained; and (iv) commonly used cutoffs such as 1 CPM might not be stringent enough to ensure a high level of reproducibility of LCPM and LFC obtained from data run at two difference sites. For example, increasing the cutoff from 1 CPM to 8 CPM increased the explained variance for fold changes from 24% to 52%.

Several groups have demonstrated that correlations are not sufficient to allow combination of datasets generated at distinct sites (16, 29). Thus, we extended our comparison to further investigate to what extent DEGs detected from Site 1 and Site 2 datasets were congruent after applying a >8 CPM low expressed gene cutoff and how gene filtering, fold change, and FDR cutoffs modulated this agreement. We found that when using >8 CPM, FDR <0.05, and FC ≥1.5 that (i) agreement increased with the strength of the biological signal (55-73% of DEGs agreeing between Days 1-14 with an average of 65% agreement) and (ii) the largest negative impact on inter-site DEG agreement was driven by genes meeting the FDR but not meeting the FC for one site but not the other (on average 68% of DEGs among genes with >8 CPM in both laboratories). These findings highlighted the importance of the accuracy of fold changes for inter-site DEG agreement. Based on these findings, our recommendation for improving inter-site DEG agreement is to apply gene expression filtering and determine DEGs using a FDR cutoff, but to not use fold-change cutoffs. In our case, after removing the fold-change cutoff requirement, the agreement in DEGs increased from 65% to 82% on average between the two sites.

In addition to the aforementioned differences, global PCA and MDS analysis of LCPM across all 100 samples showed a strong separation of the two sites indicating that the data could not simply be merged, e.g., by using the sum of the read counts obtained for the two aliquots belonging to the same sample (**Supplemental Figure 6)**. This effect could not be removed using TMM normalization, a common approach used for normalizing systematic differences in RNA-Seq data. While sequencing technology was the same, each site utilized a similar, but distinct library preparation method: Site 1 used the Illumina TruSeq method and 500 ng of RNA as input, whereas Site 2 used a modified, in house method similar to TruSeq and 1 ug of RNA for starting material. Thus, even when using highly similar library preparation methodologies, the random variation added by different sequencing laboratories is sufficient to discourage pooling the data. In addition, biological variation between aliquots derived from the same samples may explain some of these differences as well. These results have important implications for RNA-Seq experimental design considerations in the setting of large consortia, in which hundreds or thousands of samples are sequenced across multiple sites. These data indicate that, even under scenarios in which sequencing laboratories have harmonized library prep methodology, strategies that pool data at the readcount level directly should be avoided.However, biological conclusions drawn from each individual dataset on the DEG level were very similar with the heatmap fold-change profiles showing clustering of all DEGs within aliquots from the same sample for all timepoints (**Supplemental Figure 7**). This indicated that, while differences were observed on the global gene expression level, these did not sufficiently confound the key signals in the data that were accurately captured in either experiment.

### Technical Parameters

#### Choice of read length

We assessed the impact of differing read length on our ability to detect DEGs by truncating reads to mimic common Illumina sequencing configurations (100 nt SE, 75 nt SE, and 50 nt SE), and compared DEGs recovered to those found using 151 nt. Overall, for the strongest biological signal observed at Days 2-14, we found very little impact, the overlap in DEGs was >90% for all configurations, and it was 95% and 94% for the 100 nt and 75 nt simulations, respectively, based on Jaccard index. Our findings are largely in agreement with prior work examining this relationship: Chhangawala et al (17), utilized commercial reference RNA samples (UHRR and Ambion FirstChoice Human Brain Reference RNA) and found that very little difference in the sensitivity to detect DEGs when varying read length from 50-100 nt, but read lengths of 25 nt were inferior. In this regard, our study replicates and extends this principle to “real-world” samples from a vaccine trial. In contrast, recent data has emerged that indicates that short-paired end reads (i.e. 2×40 nt) outperform longer single-ended libraries for gene expression estimates (32). These findings are intriguing and warrant future investigation. However, it should be noted that the primary metrics for this determination were correlations of read counts to the “parent” dataset which comprised of 2 × 125 paired end reads. Given the size of our dataset, we did not utilize paired-end sequencing and are unable to directly test these findings. Collectively, our data demonstrate that RNA-Seq studies with clinical PBMC datasets can reduce read length down to 50 nt with relatively minimal impact on the ability to detect DEGs compared to the untruncated data. While our observations on the relatively modest impact of read count are generally consistent with prior observations, our simulation was performed using a dataset with relatively deep coverage (∼35 M reads per sample), and the impact may be more evident on datasets with less sequencing depth. Nevertheless, these results indicate that reducing read length can be a viable option for cost-savings.

#### Choice of Coverage

For this analysis, we assessed the relative impact of reducing coverage from 35 to 25, 20, 15, and 10 M reads on DEG agreement. Our main findings were that (i) compared to read length when coverage was kept constant (35 M reads), coverage had a larger influence on DEG agreement when read length was kept constant (151 nt); (ii) the relationship was modulated by the strength of the biological signal; (iii) for the strongest biological signals (Day 7 and 14), an approximate linear relationship was observed with a 5% drop in agreement with each drop in 5 M read coverage (down to 80% agreement with 10 M reads); and (iv) for the weaker signal at Day 2, a non-linear relationship was observed with the biggest drop seen between 15 M and 10 M, indicating that weaker biological signals require higher coverage than stronger signals to produce robust DEG results.

In order to assess the impact of coverage on accurately covering genes expressed in the PBMC transcriptome, based on pooled data, we generated a “PBMC transcriptome compendium” comprised of 26,172 genes. Cumulatively, our results demonstrated that, even at relatively deep sequencing coverage of 30 M, a significant percentage of the gene content expressed in PBMCs was not detectable (17-44% depending on read count filtering). This limitation in turn negatively impacts DEG detection contributing to an increased false negative rate.

#### Choice of lowly-expressed gene cutoff

Our analysis showed that stronger filtering improved the inter-site agreement – in particular, the correlation between fold-change estimates – by removing lowly-expressed genes. This prompted us to investigate the relationship between known fold changes using endogenous reference RNA (ERCC1 and ERCC2 controls) which were spiked into clinical samples in a way to evaluate fold-change accuracy (ERCC1 was spiked into 10 samples, and ERCC2 was spiked into the remaining 40 samples). Based on the observed non-linear trend, we devised a method to empirically determine a low gene expression LCPM cutoff that represents good Spearman correlation of observed vs expected ERCC control fold changes. Using this approach with each site’s data separately, we established low gene expression thresholds of 3 LCPM and 2 LCPM for Sites 1 and 2, respectively. These thresholds balance the trade-off between fold-change accuracy and filtering out too many genes for each site. These results were encouraging as they demonstrated that both sites retained a high level of technical precision to accurately quantify ERCC1/2 spike-in expression changes at relatively low CPM. However, differences remained with Site 1 requiring stronger filtering than Site 2 due to slightly lower depth of coverage and increased noise (one globally outlying sample was identified) for Site 1. This highlights the usefulness of internal ERRC1/2 spike-ins for estimating empirical gene expression filtering cutoffs accounting for systematic experimental differences across sites.

### Impact of Sample Size & Sequencing Depth on Statistical Power

One of the most important decisions to be made during the design of an RNA-Seq experiment is the number of samples to sequence. A set of “truly differentially expressed genes” or TDEGs was established based on a stringent FDR threshold and replication in both sites using different DEG detection methods. This strategy to define a “truth set” has been used effectively by prior studies examining the relationship of RNA-Seq parameters with power (33). We defined statistical power, or sensitivity, as our ability to detect TDEGs with a given effect size at varying read depths and sample sizes for each post-vaccination day. The results unambiguously demonstrated that sample size was a much more important factor than sequencing depth for detection of TDEGs in PBMCs after vaccination. A few other principles were identified:

#### (i) the power to detect gene expression with variable sample sizes was highly influenced by the effect size (i.e. the fold change)

When we restricted our analyses to genes with a greater than 2-fold change on Day 14, the power was 66% to 68% for all coverage levels considered with only n=3 samples, and this increased to >90% with n=8 samples. In contrast, when we lowered the threshold to 1.5-fold changes, the power to detect TDEGs fell to below 38% with n=3, and was below 79% with n=8. While we did not run a simulation using all genes, our simulation using a fold-change threshold of ≥1.25-fold contained between 70% to 88% of all DEGs found at each day at an FDR <0.05, and we reasoned that this was an acceptable threshold likely to screen out unreliable transcripts with extremely low effect sizes and allow us to estimate the power to detect most (if not all) of the TDEGs. Nevertheless, using the fold-change cutoff of ≥1.25-fold, even at a simulated sample size of n=8, we were only able to detect fewer than 58% of the TDEGs on Day 14. This relationship between effect size and power has been observed by others: Hart et al. found using simulated RNA-Seq datasets from PBMCs that genes with fold changes approaching 3 are nearly universally detected, but as the fold change becomes lower than 2-fold, sensitivity decreases sharply (34).

#### (ii) the sample size needed to sensitively detect the full complement of differentially expressed genes (i.e., all TDEGs at all fold changes) is remarkably high

Even with enforcing a ≥1.25-fold-change cutoff, our simulations indicated that a sample size of n=10 provided only 29% power to detect TDEGs on Day 1 and peaked on Day 14 with 68% power. This observation has been reported several times using a wide variety of datasets with differing samples: Wu et al. demonstrated that even using extremely high read depth, sample sizes of >10 were needed to achieve sensitivities above 80%. Ching and colleagues examined a diverse array of datasets: whereas some datasets were able to achieve a high level of power at low replicates (i.e., n=3 to 5), these tended to be unique scenarios such as comparing reference samples from highly divergent tissues and employing technical replicates, where average fold changes of DEGs were high, and the gene-level dispersion extremely low (42). In contrast, our work demonstrated that when analyzing human samples from diverse individuals that mimic true biological variability and comparing responses in the same tissue induced by exogenous stimuli, the sample size to achieve power >80% across all fold-change cutoffs was n≥15 for the strongest biological signal observed at Day 14. At Day 7 and Day 2, power was ≥70% followed by Day 1 with ≥50%. Schurch et al.’s findings suggest that sample sizes should be larger than n=12 if the goal is to sensitively describe the majority of DEGs (20). Hart and colleagues demonstrated that the gene-level dispersion was the main driving factor determining the sensitivity of an RNA-Seq dataset and that while datasets with low coefficients of variation could achieve 80% power at n=8, other datasets needed n>20 or even n>40. In the context of these prior studies, our power simulations illustrate the utility and limitations of RNA-Seq data using PBMCs in clinical studies. First, in our PBMC dataset, we found that at a relatively low fold-change cutoff of ≥1.5, we achieved 80% power at n=9 for TDEGs on Day 14. At this sample size, we had extremely high sensitivity to detect the genes most dramatically influenced by the perturbation. Conversely, when the gene set is defined with the fold-change threshold of ≥1.25, we estimated approximately 62% power at n=9 and attained 80% power only at n>14. In practical terms, these results provide guidance on how RNA-Seq data can be interpreted from studies using similar data. Importantly, these results do not imply that datasets where the sample size is <10 are totally unreliable. Rather, we conclude that, at smaller sample sizes, if one employs a higher fold-change cutoff, then there is a higher degree of confidence to detect true DEGs. However, more caution must be placed on interpreting genes that do not pass the FDR or fold-change threshold as true negatives – the study may be underpowered to detect them. If the goal of the experiment is to catalogue all (or nearly all) DEGs induced by a perturbation, then a large sample size is needed. While applying a fold-change cutoff increased statistical power, we have shown that inter-site agreement in DEGs was reduced when using ≥1.5 due to variation in fold-change estimates between sites. Thus, the ability to detect true TDEGs would need to be balanced with the ability to reproduce findings between different experiments.

#### (iii) the impact of sequencing depth on the power to detect TDEGs was minimal

The ability to detect TDEG’s was indistinguishable ranging from 20 M reads to 60 M reads (at 10 M read intervals). A minor loss of sensitivity was observed when samples were subsampled down to 10 M reads. This result demonstrated that a sequencing depth of 20 M reads is sufficient to capture the majority of TDEGs in a PBMCs derived dataset. Ching et al reported nearly identical findings, that minimal increased sensitivity was gained above 20 M reads (33). Similarly, others have simulated the increase in power at increased depths and found that at 2-fold depth, the increase in power only increased 3-5%, and even at 10-fold coverage, that power only increased by 2-3%.

While sequencing depth did not profoundly influence statistical power, we did observe that reducing depth had a stronger influence on transcript representation. We found that even at relatively deep sequencing depths, only approximately 65% of the total fraction of PBMC transcripts are reported in any individual sample. Decreasing sequencing depths led to a loss of content of approximately 5% of identifiable genes for each 10 M drop in depth. For example, reducing coverage from 30 M reads to 10 M reads detected 2,515 fewer genes. Taken together with our power simulations, this analysis demonstrates that while read depth does not alter overall power, a substantial proportion of genes with lower expression will be lost or provide unreliable quantification. The true impact on the biological interpretation of reduced transcript representation needs to be evaluated for each RNA-Seq dataset independently. In summary, these results demonstrated that while it is possible to reduce costs in an RNA-Seq experiment with an overall loss of power by decreasing coverage, this step must be carefully considered as there is a tangible loss of gene content. As the cost-per-base continues to drop as sequencing technology improves, the relative savings in reducing sequencing depth are becoming minimal compared to the cost of gene-content loss in an RNA-Seq experiment.

#### (iv) the false discovery rate and type-I error was dependent on the biological signal strength

Day 14 produced the strongest biological signal, having a relatively high number of TDEGs and magnitude of fold changes compared to other days. The FDR was near or below the 5% level on Day 14, regardless of the sample size, coverage level, or fold-change threshold used. As we moved back toward Day 1, the strength of the biological signal weakened at each preceding time point, and the control of the FDR became more dependent on sample size. On Day 2 and Day 1, the sample size was the most important factor for ensuring the FDR was not inflated. As with power, the sequencing depth did not have an effect on FDR as long as the depth was at or above 20 M.

The strength of the biological signal appeared to be the greatest influencer on type-I error, but this effect was balanced by the power and FDR. For example, Day 14 shows the highest type-I-error rates (although still below 2% in all cases), but this is accompanied by the highest power and lowest FDR. Sequencing depth had negligible impact on the type-I-error rate, consistent with the findings for power and FDR.

### Conclusions

In this study, our goal was to establish parameters and guidelines for RNA-Seq studies using patient PBMC samples obtained before and after vaccination with a live attenuated tularemia vaccine. Estimating statistical power in RNA-Seq studies is a complex endeavor, as power is strongly influenced by the biological variance of the particular experiment, range of effect sizes, and the technical factors we have described in this study. The near universal usage of Illumina sequencing has partially standardized the technical factors influencing RNA-Seq power. A number of recent studies have established a framework of statistical power and quality control for RNA-Seq experiments, and their findings are largely in agreement with our data (14, 16, 18). Here, we have extended these framework studies by demonstrating that the principles established hold true using “real world” clinical samples from a vaccine clinical trial. The main conclusions are: (i) filtering lowly-expressed genes is recommended, if possible guided via ERCC spike-ins, to improve fold-change accuracy and inter-site agreement – we provided an algorithm to accomplish this; (ii) read length did not have a major impact on DEG detection, and although shortening reads will result in some lost DEGs compared to longer reads, it may be a good option to save costs (iii) applying fold-change cutoffs for DEG detection reduced inter-set DEG agreement and should be used with caution, if at all; (iv) reduction in coverage had a minimal impact on statistical power but reduced the identifiable fraction of the PBMC transcriptome – 20 M reads appear to be sufficient; (v) sample size is the most important driver of statistical power followed by effect size (i.e. the magnitude of fold change) – a sample size of n=15 captured TDEGs with ≥ 1.25 fold change at > 80% power for the strongest signal (Day 14). As transcriptomics is being increasingly included in clinical drug and vaccine trials to understand biological molecular mechanisms and to provide more sensitive monitoring for toxicities, this study should provide useful guidelines for the design of multicenter RNA-Seq studies.

## Materials and Methods

Total RNA was extracted using the RNeasy kit (Qiagen, CA) and quantitated on a Thermo Nanodrop, quality assessment was performed on an Agilent Bioanalyzer. RNA with RIN score >7.5 were used for library construction.

### RNA-Seq Library Preparation and Sequencing: Site 1

Libraries were prepared using the Illumina (Illumina Inc. San Diego, CA, USA) TruSeq™ mRNA stranded kit as per manufacturer’s instructions. 500 ng of total RNA was used for library preparation. ERCC synthetic spike-in 1 or 2 (Thermo Fisher Scientific Inc., Waltham, Massachusetts) was added to each total RNA sample and processed in parallel. The TruSeq method (high-throughput protocol) employs two rounds of poly-A based mRNA enrichment using oligo-dT magnetic beads followed by mRNA fragmentation (120-200 bp) using cations at high temperature. First and second strand cDNA synthesis was performed followed by end repair of the blunt cDNA ends. One single “A” base was added at the 3’ end of the cDNA followed by ligation of the barcoded adapter unique to each sample. The adapter-ligated libraries were then enriched using PCR amplification. The amplified library was validated using a DNA tape on the Agilent 4200 TapeStation and quantified using fluorescence-based method. The libraries were normalized and pooled and clustered on the HiSeq3000/4000 flow cell on the Illumina cBot. The clustered flowcell was sequenced on the Illumina HiSeq3000 system employing a single-end 101 cycles run, each samples was sequenced to an average depth of 25 M reads.

### RNA-Seq Library Preparation and Sequencing: Site 2

Total RNA integrity was determined using Agilent Bioanalyzer or 4200 Tapestation. ERCC synthetic spike-in 1 or 2 (Thermo Fisher Scientific Inc., Waltham, Massachusetts) was added to each total RNA sample and processed in parallel. Library preparation was performed with 1 ug,of total RNA with a Bioanalyzer RIN score greater than 8.0. Ribosomal RNA was removed by poly-A selection using Oligo-dT beads (mRNA Direct kit, Life Technologies. In cases, where samples had less than 1ug of total RNA, the entire available amount was used. mRNA was then fragmented in buffer containing 40 mM Tris Acetate pH 8.2, 100 mM Potassium Acetate and 30 mM Magnesium Acetate and heating to 94 degrees for 150 seconds. mRNA was reverse transcribed to yield cDNA using SuperScript III RT enzyme (Life Technologies, per manufacturer’s instructions) and random hexamers. A second strand reaction was performed to yield ds-cDNA. cDNA was blunt ended, had an A base added to the 3’ ends, and then had Illumina sequencing adapters ligated to the ends. Ligated fragments were then amplified for 14 cycles using primers incorporating unique index tags. Fragments were sequenced on an Illumina HiSeq 3000 using single end reads extending 100 bases to an average of 35 M reads per sample.

### RNA-Seq Data Processing and Detection of DEGs

The latest version of the human reference genome (GRCh38), gene models, and associated gene annotation information at the time of study start were obtained from the ENSEMBL database (Version 84, March 2016). The genomic reference was built by merging all human chromosomes except X and Y chromosomes to avoid gender-specific effects. Reads were mapped against the human reference genome using the STAR splice-aware read aligner (Version 2.5.2a) (35). Gene expression quantification was carried out using the featureCounts function as implemented in the Subread software (Version 1.5.0-p2)(36). TMM normalization was executed as implemented in *edgeR* (37).

Genes with expression levels ≤ 8 CPM for all 50 samples within site were filtered out and were excluded in the differential gene analysis. To identify genes that were significantly differentially expressed (DE) from baseline for each post-vaccination day (Days 1, 2, 7, 14), a negative binomial model was fit to read counts using the implementation provided by the *edgeR* software(37). Each model included fixed effects for subject to account for paired samples and study visit day (baseline or post-vaccination day). For each gene, the statistical significance of the study visit day effect was evaluated using a likelihood ratio test. To control for testing multiple genes, the false-discovery rate (FDR) based on the Benjamini-Hochberg procedure as implemented in the p.adjust R function was applied for each model (38). Genes with a fold change from baseline ≥ 1.5 and FDR-adjusted p-value < 0.05 were considered DEGs.

### Simulation of datasets to assess impact of read coverage, read length, and simulate power

The original 50 FASTQ files from Site 1 with a read length of 151 nt were right-truncated to obtain three additional FASTQ files per sample with 50 nt, 75 nt, and 100 nt long reads. Truncation of reads was carried out using the fastx_trimmer tool of the FASTX Toolkit. A set of four FASTQ files per sample to simulate different coverage levels were obtained by randomly down-sampling original reads to 25 million (25 M), 20 million (20 M), 15 million (15 M), and 10 million (10 M) reads. Random subsampling of reads was carried out using the sample tool of the seqtk Toolkit (Version 1.2-r95-dirty). An additional four FASTQ files were simulated for the power analysis using the same approach, which contained 30 M, 40 M, 50 M, and 60 M reads, respectively.

### Empirical determination of a cutoff for filtering lowly-expressed genes using ERCC controls

The 92 ERCC 1 mix transcripts were added to pre -vaccination samples, and the ERCC 2 mix was added to each post-vaccination sample. ERCC spike ins cover a range of concentrations from 0.014 to 30,000 CPM for both mixes, and the expected LFCs from mix 2 to mix 1 range from 0.25 to 2. For paired samples, the empirical fold change (based on fold change in CPM from the RNA-Seq experiment) and expected fold changes were determined for each of the 92 ERCC transcripts. Likewise, the mean LCPM of each ERCC transcript was determined by first calculating the average LCPM value for each paired sample, and then taking the mean of those averages. This results in a weighted average for each ERCC transcript across the pre- and post-vaccination samples. Based on the distribution of the mean ERCC LCPM, the 92 spike-ins were allocated into 7 bins of increasing abundance from low to high (≥13 transcripts per bin).

Next, for each paired sample, the Spearman correlation between the empirical and the expected fold changes was calculated for the subset of ERCC transcripts within each bin. The mean LCPM of the transcripts within each bin was also computed for each paired sample. This produces a pair of values – the Spearman correlation and mean LCPM – for each paired sample within each bin of ERCC transcripts. A 3^rd^ order polynomial was fitted to these correlation data to estimate a trend line that captured the non-linear relationship between the correlation and average LCPM. To make this step more robust, outliers within each bin (points with correlation values exceeding Q3 + 1.5 IQR or less than Q1 – 1.5 IQR) were exclude from this step. The LCPM cutoff value was then chosen as the value at which the fitted polynomial gave a correlation of 0.9. Bootstrapping (sampling with replacement of paired samples) was used to determine the 95% confidence interval of the cutoff point; the lower and upper bound of the confidence interval represented the 2.5 and 97.5 percentiles, respectively, of the bootstrap cutoff values.

### Calculation of Statistical Power

Statistical power was assessed using a modified version of the PROPER (39, 40) R package. The simulation strategy was designed to assess the effect of (i) the coverage level of the RNA-Seq experiment, (ii) the log fold-change cutoff used to filter genes, (iii) the sample size, and iv) the varying strength of the biological signal post-vaccination (Day 1, 2, 7, and 14).

For each post-vaccination day, we defined a gold standard set of genes that were “truly differentially expressed genes” (TDEG) as those genes with maximum expression level of at least 8 CPM for all 50 samples within a site and an FDR-adjusted p-value below 0.05, independently detected in the data from both sites by both the edgeR (37) and DESeq2 R packages (41), using the original coverage data (Table 1). Using these criteria, these analyses yielded 444, 357, 2,300, and 3,284 TDEGs for post-vaccination Days 1, 2, 7 and 14 respectively. Each TDEG was associated with an effect size based on the estimated average fold change (FC) for that gene in the original dataset from Site 1. All non-TDEGs were associated with a null effect size (FC=1).

We used the most complete set (Site 2) that did not have any outliers for the power assessment. To estimate statistical power to detect TDEGs, different coverage levels were simulated by down-sampling the FASTQ files from Site 2 to 30 M, 20 M, 10 M reads and over-sampling them to sizes of 40 M, 50 M, and 60 M reads. We simulated FASTQ files for each of these different coverage levels for each subject and each of the five time points (pre-vaccination and each post-vaccination day). The simulated FASTQ files were aligned against the reference and processed to obtain corresponding gene count datasets as described for the original data. The same genes that were filtered out in the original analysis were also removed from these datasets. Finally, the average gene expression levels (normalized average LCPM) and dispersion estimates (trended dispersion) were calculated based on the pre-vaccination day data, resulting in paired mean and dispersion estimates for each gene and for each coverage level.

These estimates were then used to simulate gene count datasets for the power assessment: for each coverage level, post-vaccination day, and sample size (*n*), 500 datasets were simulated using independent negative binomial models for each gene parameterized by its estimated mean expression, dispersion, and effect-size level. In each dataset, *n* samples were simulated for both the pre- and post-vaccination day. The TDEGs were simulated so that the average difference in expression between the pre- and post-vaccination days was equal to its estimated effect size. The model did not include a dependence between paired samples. The simulated dataset was then analyzed with edgeR to identify differentially expressed genes. The estimated FC and p-value for each gene were calculated and recorded.

The statistical power for each combination of parameters was determined as the average proportion of TDEGs that were identified in the simulated datasets with an FDR-adjusted p-value <0.05. Four levels of fold-change cutoffs were considered: 1.25, 1.5, 1.75, and 2.0. The power calculation was conditioned on the given fold-change criteria, which means the TDEGs considered had an effect size above the cutoff value.

## Data and materials availability

The RNA-Seq data has been deposited in NCBI GEO (GSE149809).

## Ethical Statement

Subjects gave their consent for inclusion before they enrolled in the parent clinical trial (NCT01150695) (30). The protocol and consent form were reviewed by the US Food and Drug Administration, and approved and monitored by the clinical sites’ institutional review boards.

## Acknowledgements

This study was funded through the Vaccine Treatment and Evaluation Units (VTEU) awarded to Emory University (HHSN272201300018I) and St. Louis University (HHSN27220130021I). We thank the Emory VTEU administrative and finance core for their support including Dean Kleinhenz, Hannah Huston, and Michele Paine McCullough. The Emory NPRC Genomics Core is supported in part by NIH P51 OD011132 and S10 OD026799. We also thank the Georgia Research Alliance.

## SUPPLEMENTAL TABLES

**Supplemental Table 1:** Impact of fold change and FDR-adjusted p-value cutoff on DEG agreement. Total represents reference DEGs. All other columns to the right refer to the comparison group. FC = fold change ≥1.5; FDR = FDR-adjusted p-value <0.05.

| Reference Laboratory | Comparison Laboratory | Day | DE Total | Missing (% Total) | DE (% Total) | Not DE (% Total) | Fail FC and FDR (% Not DE) | Pass FC and Fail FDR (% Not DE) | Fail FC and Pass FDR (% Not DE) |
| --- | --- | --- | --- | --- | --- | --- | --- | --- | --- |
| Site 1 | Site 2 | Day 1 | 165 | 8 (4.8%) | 90 (54.5%) | 67 (40.6%) | 24 (35.8%) | 2 (3%) | 41 (61.2%) |
| Site 1 | Site 2 | Day 2 | 213 | 10 (4.7%) | 145 (68.1%) | 58 (27.2%) | 23 (39.7%) | 1 (1.7%) | 34 (58.6%) |
| Site 1 | Site 2 | Day 7 | 607 | 41 (6.8%) | 407 (67.1%) | 159 (26.2%) | 20 (12.6%) | 2 (1.3%) | 137 (86.2%) |
| Site 1 | Site 2 | Day 14 | 1086 | 72 (6.6%) | 794 (73.1%) | 220 (20.3%) | 14 (6.4%) | 7 (3.2%) | 199 (90.5%) |
| Site 2 | Site 1 | Day 1 | 155 | 26 (16.8%) | 90 (58.1%) | 39 (25.2%) | 14 (35.9%) | 9 (23.1%) | 16 (41%) |
| Site 2 | Site 1 | Day 2 | 240 | 25 (10.4%) | 145 (60.4%) | 70 (29.2%) | 32 (45.7%) | 6 (8.6%) | 32 (45.7%) |
| Site 2 | Site 1 | Day 7 | 576 | 68 (11.8%) | 407 (70.7%) | 101 (17.5%) | 21 (20.8%) | 6 (5.9%) | 74 (73.3%) |
| Site 2 | Site 1 | Day 14 | 1087 | 119 (10.9%) | 794 (73%) | 174 (16%) | 23 (13.2%) | 4 (2.3%) | 147 (84.5%) |

## SUPPLEMENTAL FIGURES

**Supplemental Figure 1:**
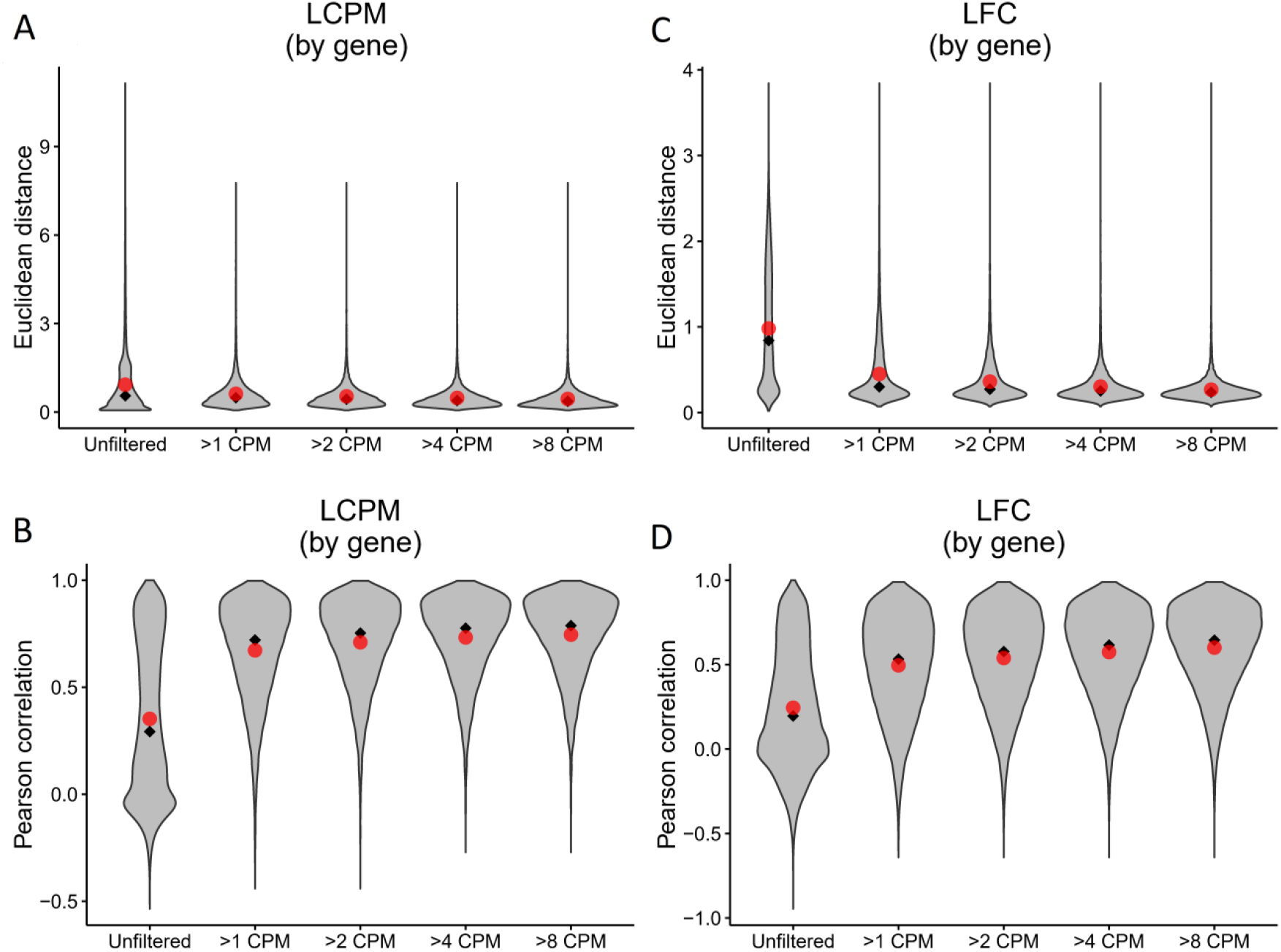
Inter-site agreement of RNA-Seq log_2_ counts per million and fold-change estimates improves after filtering out lowly-expressed genes. LCPM: log_2_ counts per million. LFC: log_2_ fold change. **(A)** Violin plot depicting the distribution of adjusted Euclidean distance between Site 1 and Site 2 of the LCPM for each gene. A total of 49 paired observations were available for each gene. Red dots mark the mean of the distribution, and black diamonds mark the median. Each column shows the distribution after filtering genes at the CPM thresholds indicated on the X axis. **(B)** The same analysis and plotting design as described in (A), but of Pearson correlation between Site 1 and Site 2 of the LCPM for each gene. **(C)** and **(D)** show the same results but calculated on the LFCs. A total of 36 paired observations of LFC values were available for each gene.

**Supplemental Figure 2:**
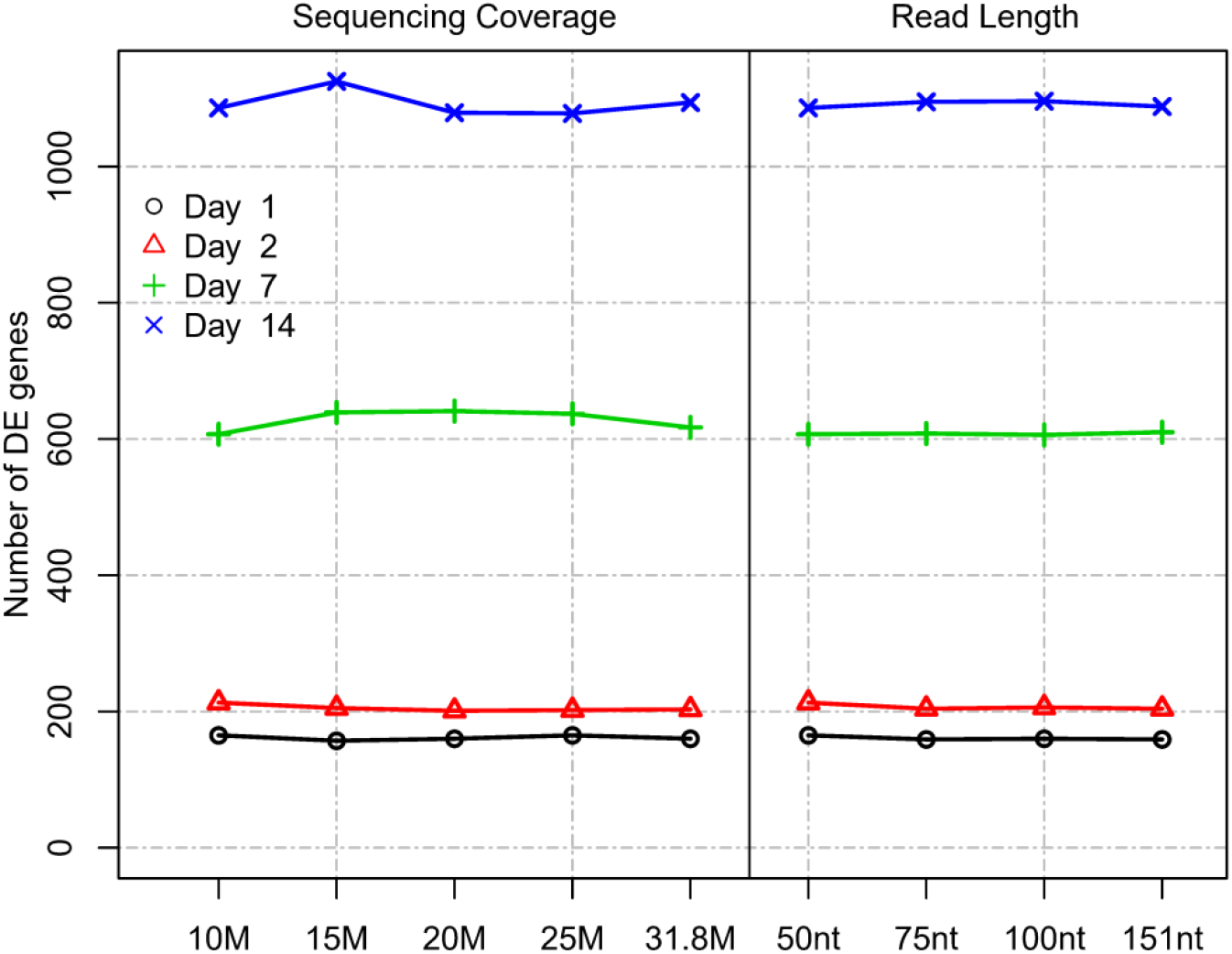
Impact of read length and coverage on DEG detection. Impact of read length and coverage on the number of DEGs (Site 1). Values for the original dataset, with a median coverage of 31.8 M and a read length of 151 nt, for comparison.

**Supplemental Figure 3:**
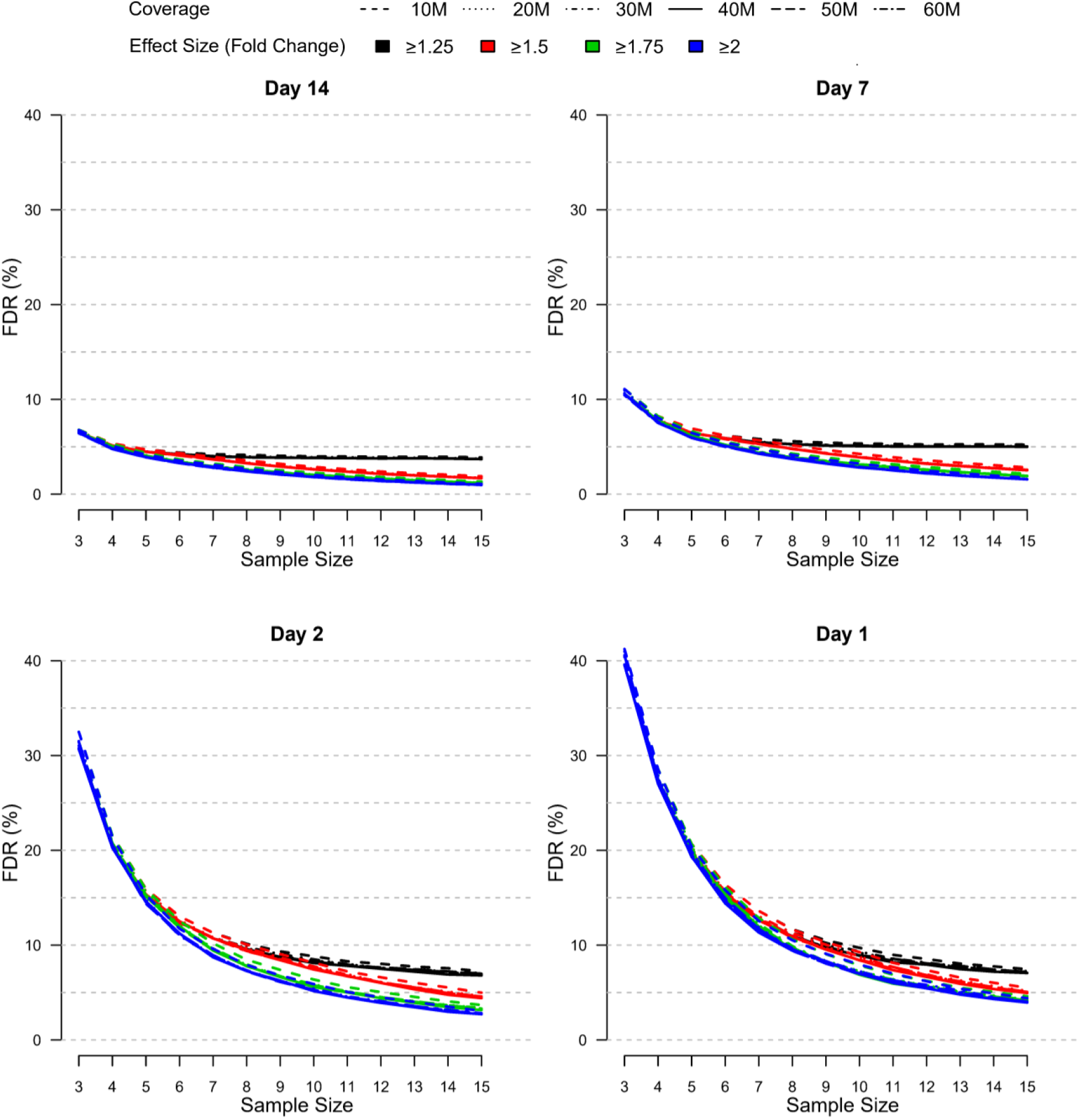
Relative false discovery rate by samples size, effect size, and coverage at each post-vaccination day as simulated using the modified PROPER R package. Days were sorted by decreasing vaccination effect based on overall fold changes and DEG responses observed for this study (see **Figure 3A**). False discovery rate was assessed for different fold-change cutoffs (indicated by color-coded lines), coverage (as indicated by the line type), and sample size (x-axis).

**Supplemental Figure 4:**
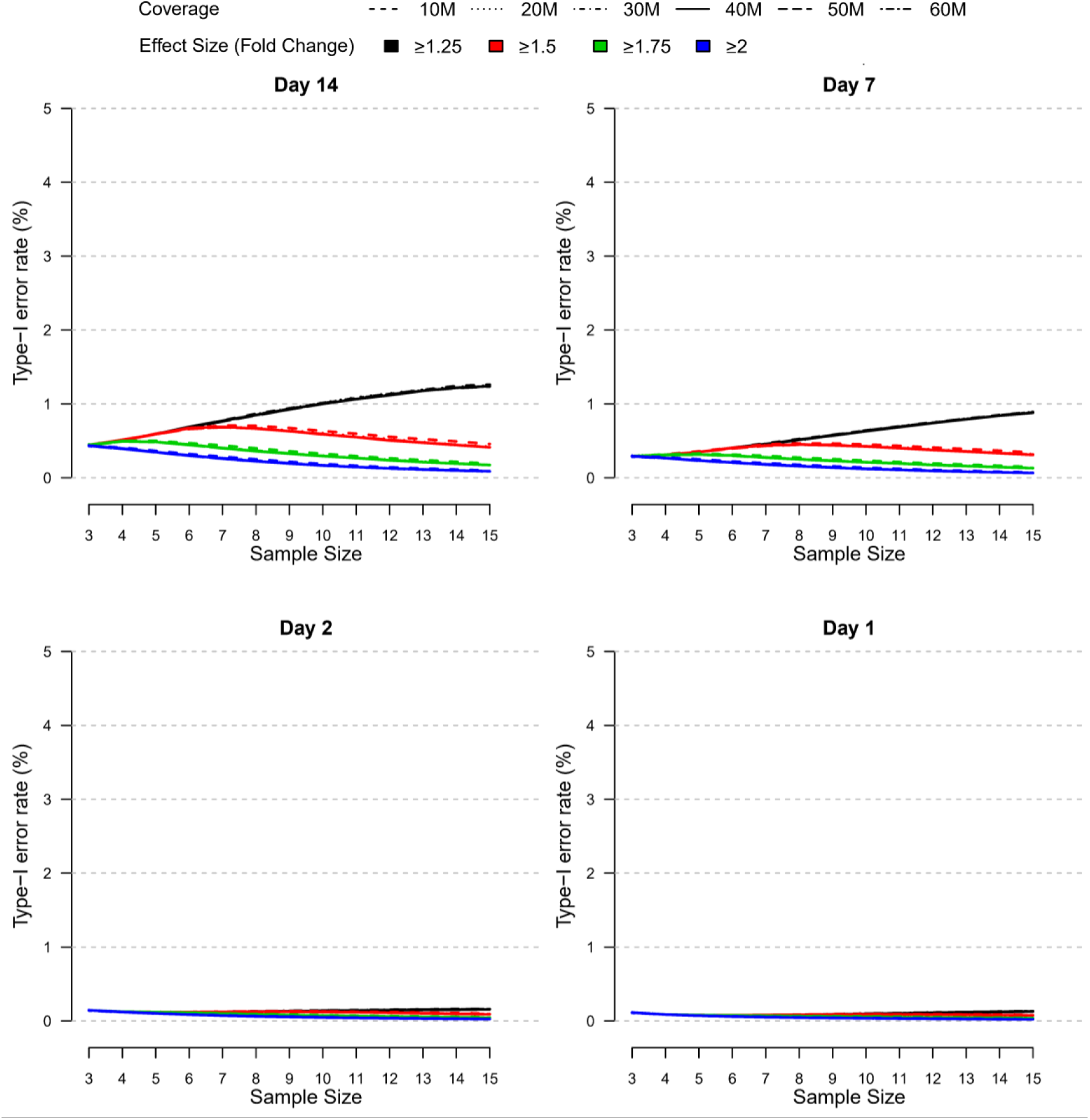
Relative type-I-error rate by samples size, effect size, and coverage at each post-vaccination day as simulated using the modified PROPER R package. Days were sorted by decreasing vaccination effect based on overall fold changes and DEG responses observed for this study (see **Figure 3A**). Type-I-error rate was assessed for different fold-change cutoffs (indicated by color-coded lines), coverage (as indicated by the line type), and sample size (x-axis).

**Supplemental Figure 5:**
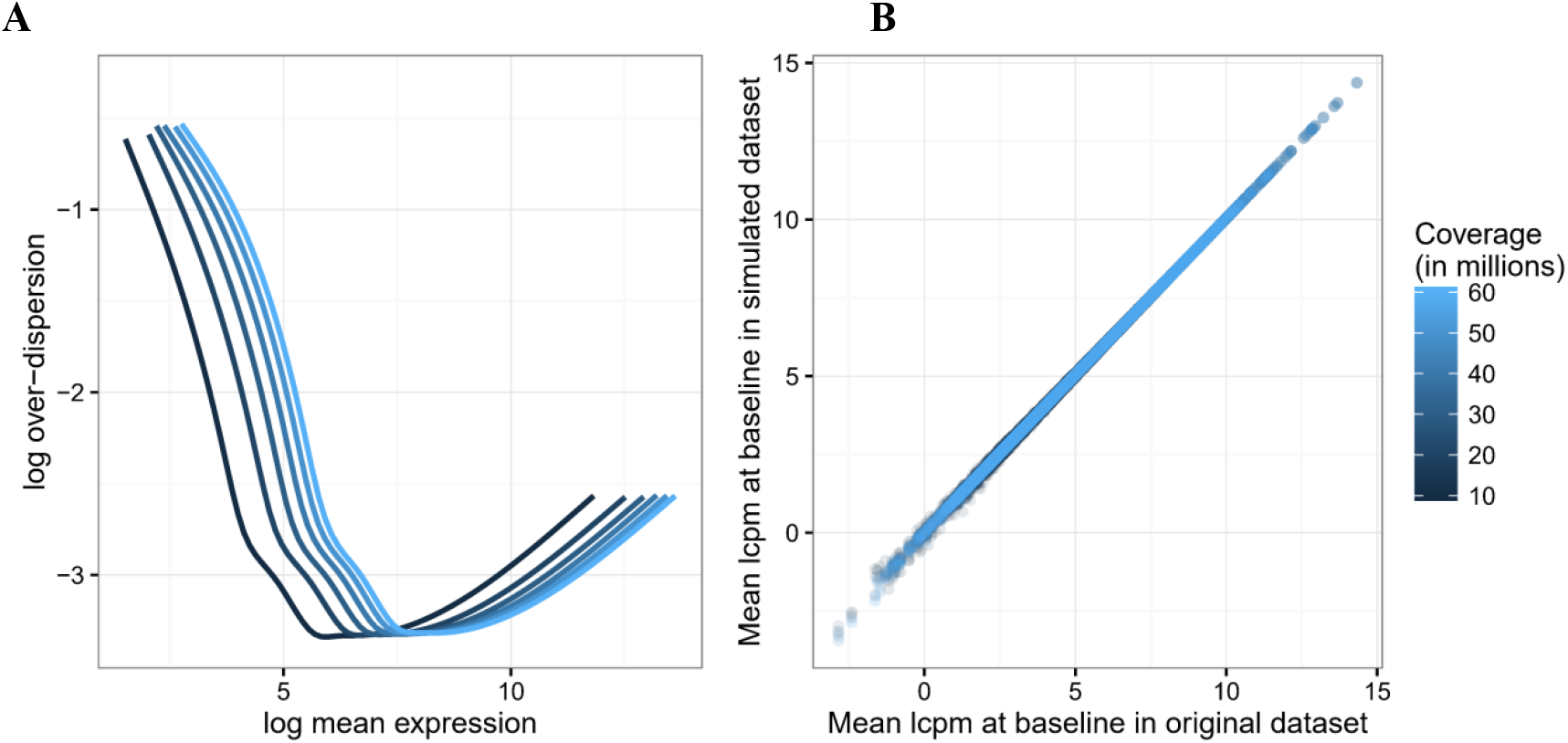
Mean-dispersion trend for each simulated coverage dataset (left) and scatterplot of mean log_2_ CPM of genes in original dataset compared to each simulated coverage dataset (right). For each gene in the simulated coverage datasets, the estimated log_2_ mean expression and log_2_ over-dispersion values were calculated; from these points, a loess curve is fit show the mean-dispersion trend at each simulated coverage level (A). The average TMM-normalized log_2_ CPM values are also calculated for each gene across baseline samples in each simulated dataset and compared to the corresponding values from the original dataset (B).

**Supplemental Figure 6:**
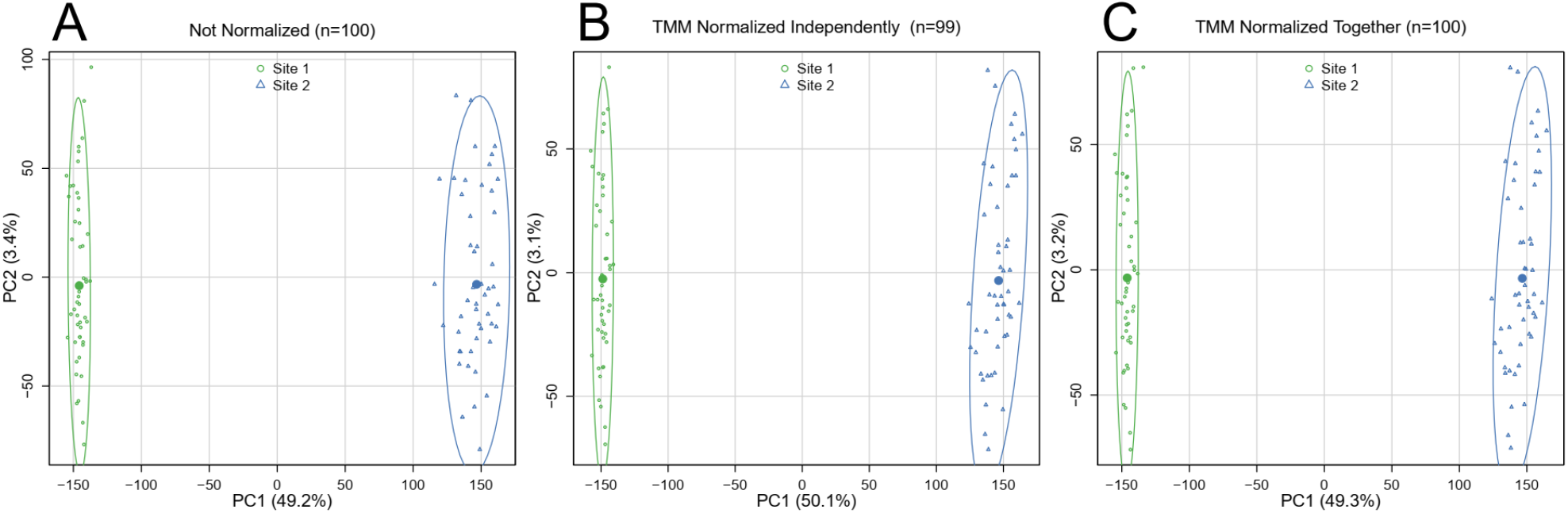
Principal component analysis of log_2_ counts per million across sites before and after normalization within and between sites.

**Supplemental Figure 7:**
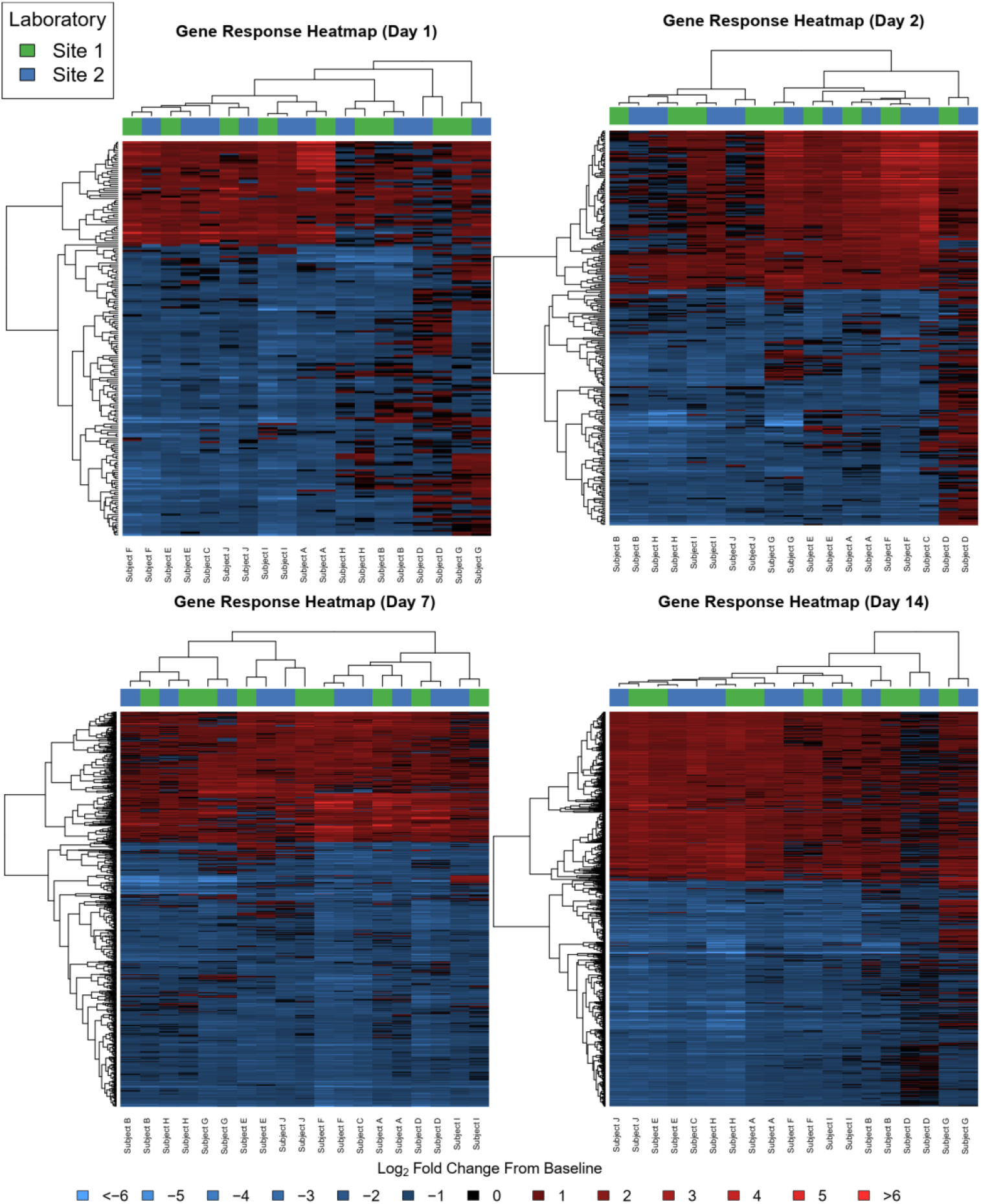
Heatmap summarizing fold change responses of the union of DEGs by subject and Site. Subjects and genes were clustered using Uncentered Pearson Correlation distance in combination with complete linkage clustering.

